# An extracellular vesicle biogenesis-inspired engineering platform for efficient protein delivery and therapeutic base editing

**DOI:** 10.64898/2026.05.18.721905

**Authors:** Songpu Xie, Qiangbing Yang, Nazma Ilahibaks, Kaiyong Qu, Bing Yao, Pieter Vader, Maike A D Brans, Christian Snijders Blok, Anders Gunnarsson, Pieter A Doevendans, Junjie Xiao, Raymond Schiffelers, Zhiyong Lei, Joost P.G. Sluijter

## Abstract

Efficient and controllable delivery of genome-editing proteins remains a central challenge for therapeutic translation of gene-editing technologies. Extracellular vesicles (EVs) offer an attractive non-viral delivery modality due to their biocompatibility and large capacity for cytosolic cargo delivery. Yet, rational strategies to achieve controlled and programmable protein loading are still lacking.

Here, we present NEO-TOP-EVs, an EV biogenesis-guided engineering platform that systematically integrates key features of three design principles inspired by vesicle formation: 1) PI(4,5)P₂-mediated plasma membrane targeting, 2) ESCRT-dependent membrane scission, and 3) self-assembly-driven cargo clustering for enabling efficient encapsulation of genome-editing ribonucleoproteins. Together, the NEO design increased cargo incorporation and enhanced functional delivery of gene editing modalities under particle-normalized conditions.

Using NEO-TOP-EVs, we achieve efficient delivery of Cas9 and adenine base editor ribonucleoproteins without nucleic acid templates. In an in vitro proof-of-concept, delivery of an adenine base editor targeting proprotein convertase subtilisin/kexin type 9 (PCSK9) induces efficient splice-site disruption, resulting in reduced PCSK9 expression and enhanced LDL receptor activity. Proof-of-concept in vivo experiments provide preliminary evidence of functional Cre protein delivery to the liver.

Together, these findings establish NEO-TOP-EVs as a modular platform for protein-based genome editing, demonstrating how biogenesis-informed EV engineering yields functional protein delivery at levels relevant to therapeutic development.

## 1. Introduction

The ability to deliver therapeutic molecules, including proteins, nucleic acids, and genome-editing complexes, into specific target cells remains a fundamental challenge for the development of next-generation therapeutics. Many disease-relevant targets are intracellular, yet current delivery strategies suffer from limited efficiency, off-target effects, immunogenicity, or poor compatibility with protein-based cargoes ^1, 2^. Therefore, there is a need for delivery vehicles that can transport functional macromolecules into cells in a controlled, safe, and programmable manner.

Extracellular vesicles (EVs) have emerged as a promising candidate to address this gap. As cell-derived natural carriers, EVs mediate intracellular communication through the transfer of proteins, nucleic acids, and lipids, and their inherent biocompatibility, low immunogenicity, and stability in physiological fluids have generated considerable interest in EVs as therapeutic delivery vehicles ^3–8^. Despite these advantages, achieving efficient and programmable loading of protein cargos, particularly CRISPR ribonucleoproteins (RNPs), remains a major barrier to translational applications.

Current EV engineering strategies, such as simple cargo overexpression ^9^, membrane permeabilization ^10^, or fusion to sorting peptides ^11^, offer limited control over encapsulation efficiency, high variability, and often yield vesicles with inconsistent cargo content ^6, 12, 13^. A generalizable engineering approach enabling effective, scalable, and controllable protein loading is therefore critically needed.

In cells, EV biogenesis is driven by a series of highly coordinated membrane remodeling events that share conserved mechanistic features across biological processes ^3^. These events include: (i) spatially confined membrane targeting to PI(4,5)P₂–rich microdomains ^14, 15^, (ii) local membrane deformation induced by protein–lipid or protein–protein interactions ^16, 17^, and (iii) membrane neck constriction and closure facilitated by ESCRT-machinery ^18–20^. These principles govern how cells package materials into intraluminal vesicles or microvesicles and thereby might offer a mechanistic foundation for rational EV engineering. To date, despite their potential, these biogenesis principles have not been systematically harnessed or integrated into a unified platform to enhance EV-mediated protein delivery.

We previously developed TOP-EVs (Technology of Protein loading through Extracellular Vesicles), an engineered EV platform that recruits cargos to budding sites via chemically induced dimerization, enabling cargo loading into released vesicles ^21^. The TOP-EV strategy enabled efficient cargo loading and functional protein delivery, demonstrating the feasibility of EV-mediated protein transport ^22^. In the original TOP-EVs design, cargo recruitment relied on N-terminal myristoylation as a generic membrane-anchoring strategy ^23, 24^. Although N-terminal myristoylation enables membrane association and effective protein delivery ^25, 26^, myristoylation does not confer intrinsic specificity for phosphoinositide or plasma-membrane microdomains ^23, 24, 27^. As a result, cargos may distribute broadly with intracellular membranes rather than concentrating at membrane domains that are competent for coordinated curvature generation, ESCRT recruitment, and vesicle scission ^18^. Because efficient EV biogenesis requires coordinated cargo recruitment and membrane remodeling ^3, 18^, this lack of spatial coupling may reduce cargo loading per vesicle and limit delivery efficiency ^13^.

To address this limitation, we developed NEO-TOP-EVs (Next Evolutionary Optimization of TOP-EVs), an EV biogenesis-guided engineering platform in which PI(4,5)P₂-mediated membrane targeting, ESCRT-dependent membrane scission, and self-assembly-driven multivalent clustering act in concert to promote coordinated vesicle formation and cargo encapsulation. As a proof-of-concept therapeutic application, we demonstrate efficient delivery of adenine base editor RNP ^28^ without the use of viral vectors or DNA templates ^29^, achieving precise therapeutic base editing of PCSK9 ^30^ and functional modulation of the LDL receptor pathway.

Together, these results establish NEO-TOP-EVs as a biogenesis-informed platform for protein delivery and highlight the importance of coordinating multiple steps of vesicle formation to enhance EV-mediated cargo loading and functional delivery.

## 2. Materials and methods

### 2.1 Cell culture

Human embryonic kidney 293FT (HEK293FT) (R70007, Thermo Fisher), T47D Cre stoplight reporter cells ^31^ kindly provided by Linglei Jiang, Cas9 stoplight reporter cells ^32^ and ABE stoplight reporter cells ^33^ kindly provided by Olivier G. de Jong, and Huh7 cells were maintained in Dulbecco’s Modified Eagle Medium (DMEM, 41965-039, Gibco) supplemented with 10% fetal bovine serum (FBS, 35-079-CV, Corning) and 1% penicillin/streptomycin (P/S, 15-140-122, Gibco). All cells were maintained at 37 °C in a humidified atmosphere with 5% CO₂.

### 2.2 Plasmid construct generation

Plasmid constructs were generated by extending the previously described TOP-EVs platform ^21^ with additional envelopment modules to create the NEO-TOP-EVs architecture. In brief, DmrA expression constructs were designed as modular fusion proteins; depending on the generation, these comprised combinations of a PI(4,5)P₂-binding domain, linker regions, an ESCRT-recruiting domain, and self-assembling peptide modules, all fused to the DmrA dimerization domain, while cargo proteins (Cre, Cas9, or ABE8e) were fused to DmrC.

Individual domains were assembled into the pcDNA3.1+ backbone using NEBuilder® HiFi DNA Assembly (E2621L, BIOKÉ) according to the manufacturer’s instructions. Detailed domain composition, orientation, and linker sequences for all constructs used in this study are provided in Figure 1 and Supplementary Tables 1-4. All plasmids were sequence-verified prior to use.

**Fig. 1.**
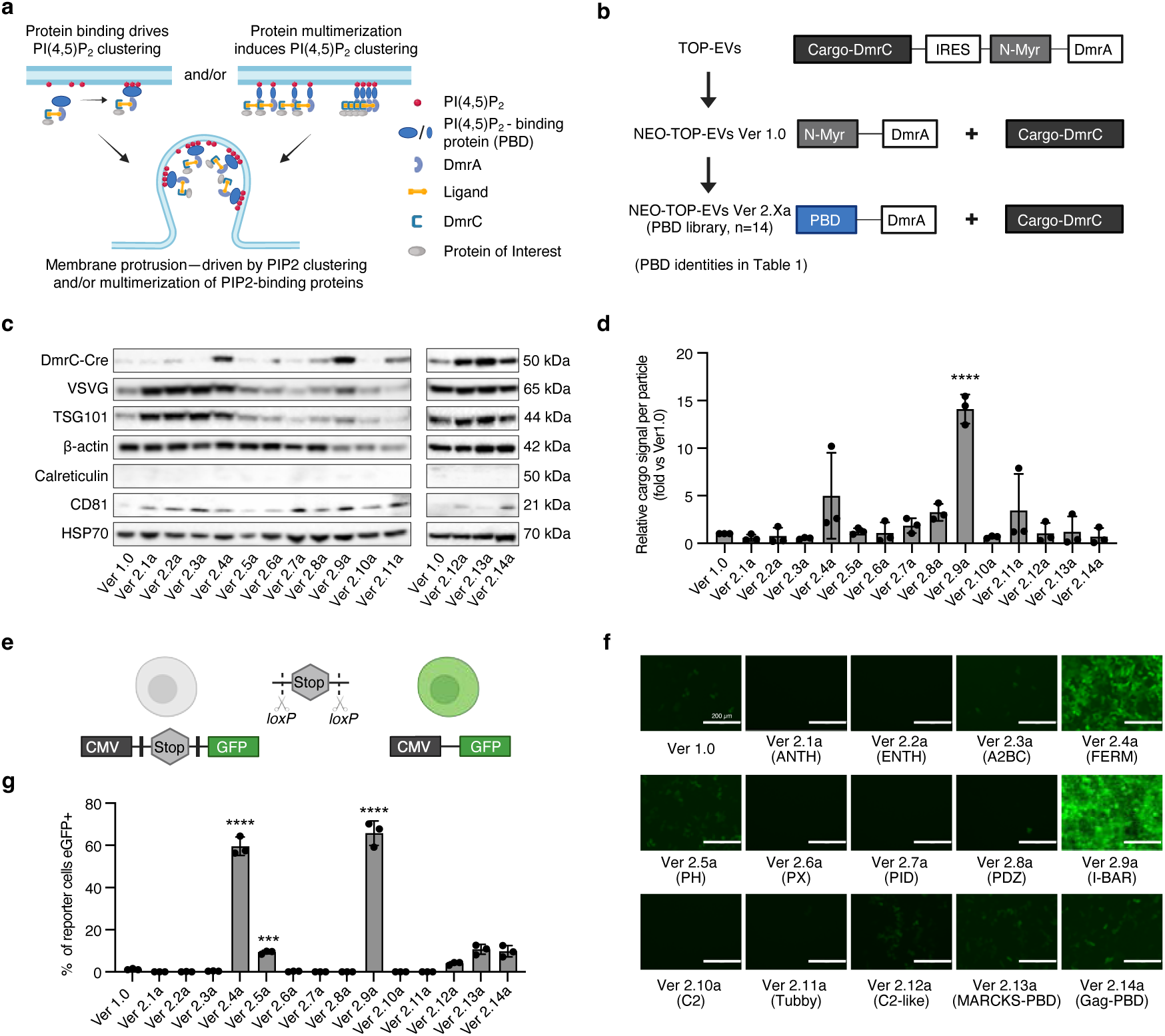
Targeting PI(4,5)P₂-rich plasma membrane microdomains enhances cargo loading and delivery efficiency. **a**, Conceptual schematic illustrating two complementary mechanisms that promote local PI(4,5)P₂ clustering at the plasma membrane: recruitment of PI(4,5)P₂-binding domains (PBDs) and multimerization-driven co-clustering of PBD-containing proteins. Local PI(4,5)P₂ enrichment and multivalent protein assembly promote membrane protrusion and vesicle budding. **b**, Design evolution from the original TOP-EVs platform to NEO-TOP-EVs. The original TOP-EVs use N-terminal myristoylation (N-Myr) for membrane anchoring and DmrA–DmrC heterodimerization for cargo tethering. In NEO-TOP-EVs Ver 1.0, a minimal N-Myr–DmrA anchor is used. In second-generation constructs (Ver 2.Xa), the N-Myr anchor is replaced by a library of PI(4,5)P₂-binding domains (PBDs). **c**, Immunoblot analysis of purified EVs produced by Ver 1.0 and second-generation NEO-TOP-EV constructs (Ver 2.1a–Ver 2.14a). Blots were probed for DmrC-Cre cargo, vesicular stomatitis virus glycoprotein (VSVG), the EV marker TSG101, β-actin, and EV-associated markers Calreticulin, CD81 and HSP70. EV were normalized to 1× 10^10^ particles per lane as determined by NTA. This normalization was applied to all EV immunoblot analyses throughout the study. **d**, Quantification of cargo incorporation per particle derived from the immunoblots shown in panel c. Densitometry values were normalized to the Ver 1.0 control within each blot. Data represent mean ± s.d. from n = 3 independent EV preparations. **e**, Schematic of the Cre stoplight reporter assay used to quantify functional EV-mediated cargo delivery. Cre recombinase– mediated excision of a loxP-flanked DsRed-stop cassette activates GFP expression, converting unrecombined reporter cells into recombined GFP-positive cells. **f**, Representative fluorescence microscopy images of T47D Cre-reporter cells treated with EVs from Ver 1.0 and Ver 2.Xa constructs. Scale bars, 200 μm. **g**, Flow cytometry quantification of GFP-positive reporter cells corresponding to panel f. Data represent mean ± s.d. from n = 3 independent experiments. For functional delivery assays, 25,000 reporter cells were treated with 1 × 10^11^ EV particles per condition, normalized by NTA. For all bar graphs, statistical comparisons versus Ver 1.0 were performed using one-way ANOVA with Dunnett’s multiple comparisons test. *** p < 0.001, **** p < 0.0001.

### 2.3 EV production and isolation

HEK293FT cells were seeded in a T175 flask and cultured overnight to reach ∼70–80% confluency prior to transfection. Cells were subsequently co-transfected with plasmids encoding the DmrC-fused cargo protein (Cre-DmrC, Cas9-DmrC or ABE8e-DmrC), the corresponding DmrA envelopment construct, and VSV-G (10 μg, 10 μg, and 5 μg per flask, respectively), using Lipofectamine® 3000 Transfection Reagent (L3000008, Invitrogen) at a DNA-to-reagent ratio of 1:2 according to the manufacturer’s instructions. After 6 hours, the medium was changed to DMEM, supplemented with 10% FBS and 1% P/S. After 24 hours, the medium was changed to DMEM supplemented with A/C Heterodimerizer (635056, Takara). After 48 h, the conditioned medium was collected and centrifuged at 500 × g for 5 min at 4 °C (repeated three times) to remove cells and large debris, followed by centrifugation at 2000 × g for 20 min at 4 °C to remove smaller cell debris and apoptotic bodies. Next, the supernatant was filtered through a 0.45 μm SCFA membrane syringe filter (516-1954, Corning). Finally, the supernatant was recovered and centrifuged at 100,000× g for 70 min at 4 °C to pellet the EVs.

### 2.4 Nanoparticle Tracking Analysis

The size and particle concentration of EVs was assessed with nanoparticle tracking analysis (NTA) (NS500, Malvern Nanosight). EVs were suspended in PBS and measured in triplicate with individual measurements of 30 seconds at camera level 15. Analysis was performed with NTA software 3.3 with a minimal track length of 10, detection threshold 5, and screen gain 1. The cut-off values for reliable measurement were between 20.0 and 100.0 particles/frame.

### 2.5 Western blotting

Cell lysates and EVs were lysed in RIPA lysis buffer (1:10, 20-188, Sigma) supplemented with Protease/Phosphatase inhibitor cocktail (1:100, 5872S, Cell Signaling Technology) and stored on ice for 30 min. Next, samples were centrifuged at 14,000 × g for 10 min. at 4 °C where the supernatant was isolated. The lysed EVs were stored at -80 °C. For EV western blot analysis, samples were normalized to 1 × 10¹⁰ particles per lane as determined by NTA prior to loading.

Protein concentrations were determined via the Micro BCA Protein Assay Kit (23235, ThermoFisher). Samples were mixed with NuPAGE™ Sample Reducing Agent (50 mM, NP0004, Invitrogen Corp) and NuPAGE LDS Sample Buffer (NP0007, Invitrogen), incubated at 70 °C for 10 min. Samples were separated on Bolt™ 4–12% Bis-Tris Plus Gel (NW04125BOX, ThermoFisher Scientific), with a PageRuler Plus Prestained Protein Ladder (26619, ThermoFisher Scientific) at 200 V for 35 min and transferred to PVDF membranes (IPVH00010, Merck). The membranes were blocked for 1 h in Intercept® (TBS) Blocking Buffer (927-60003, LICORbio).

Primary antibodies included Rabbit anti-DmrC (1:1000 dilution, Takara, 635091), Rabbit anti-TSG101 (1:1000 dilution, Abcam, ab30871), mouse anti-β-Actin (1:5000, Cell Signaling Technology, 3700s), mouse anti-Cas9 (1:1000, Cell Signaling Technology, #14697, clone 7A9-3A3), anti-rabbit Cre recombinase (1:1000, Cell Signaling Technology, #15036, clone D7L7L), anti-mouse PCSK9 (1:1000, #MA5-32843, ThermoFisher, clone 2F1), anti-rabbit LDLR (1:1000, #MA5-32075, ThermoFisher, clone SJ0197). Secondary antibodies included Goat Anti-Rabbit Immunoglobulins/HRP (1:2000, Dako, P0448) and Goat Anti-Mouse Immunoglobulins/HRP (1:1000, Dako, P0447).

### 2.6 Reporter assay

In a flat bottom 96-well plate (655075, Greiner CELLSTAR), Cre T47D stoplight reporter cells were seeded at a density of 25,000 cells/well (for Cas9/ABE stoplight reporter, cells were seeded at a density of 10,000 cells/well) and incubated overnight. The next day, the conditioned media or differential ultra-centrifuged EVs were added to T47D stoplight reporter cells. After 3 days, the cells were directly visualized with EVOS Cell Imaging System (M5000, Invitrogen) and subsequently analyzed by flow cytometry (CytoFLEX, Beckman Coulter).

### 2.7 Flow cytometry analysis

Cells were washed with PBS and dissociated with 0.25% Trypsin-EDTA solution (Sigma-Aldrich, T4049). The dissociated cells were transferred to a 96-well round bottom plate (650185, Greiner CELLSTAR ®), centrifuged at 500 × g for 3 min, and resuspended in 250 µl PBS supplemented with 2% FBS. Fluorescence data were acquired by employing a CytoFLEX flow cytometer (Beckman Coulter, Inc.) and analyzing using Kaluza software v2.1 (Beckman Coulter, Inc.). Standard forward and side scatter gating was applied to exclude debris and doublets.

For surface LDLR staining, cells were washed with PBS, dissociated with 0.25% Trypsin-EDTA, and transferred to a 96-well round bottom plate. Cells were centrifuged at 500 × g for 3 min and resuspended in PBS supplemented with 2% FBS. Cells were incubated with APC-conjugated anti-human LDLR antibody (FAB2148A, R&D Systems) at 10 µL per 10⁶ cells for 30 min at 4 °C, washed with PBS, and resuspended in PBS with 2% FBS prior to acquisition. Fluorescence data were acquired using a CytoFLEX flow cytometer and analyzed with Kaluza software v2.1. Results are expressed as mean fluorescence intensity (MFI) fold change relative to NC.

### 2.8 Microscopy analysis

Microscopy analysis was performed using the EVOS FL Cell Imaging System (Life Technologies). Live cell brightfield and fluorescence imaging was used to visualize reporter activation and assess cell morphology following EV treatment. Images were acquired using identical acquisition settings within each experiment.

### 2.9 Transmission electron microscopy (TEM)

EVs were obtained via differential ultracentrifugation and suspended in PBS. EV samples were adsorbed onto carbon-coated grids (75–200 mesh) for 15 min at room temperature. Grids were washed with PBS and fixed with 2% PFA and 0.2% glutaraldehyde in PBS for 30 min at room temperature. Grids were subsequently stained with uranyl-oxalate and embedded in 1.8% methylcellulose containing 0.4% uranyl acetate for 10 min on ice. EVs were visualized using a JEOL transmission electron microscope.

### 2.10 In vivo Cre delivery in Ai9 reporter mice

Animal care and handling were carried out in accordance with the approval of the Animal Ethical Experimentation Committee of Utrecht University, The Netherlands, and per the guidelines of the “Guide for the Care and Use of Laboratory Animals” under work protocol no. 105252-1. B6.Cg-Gt(ROSA)26Sor^tm9(CAG-tdTomato)Hze/J transgenic mice (Ai9 mice, strain #007909), obtained from The Jackson Laboratory, were kept in our local breeding facility. Ai9 mice were housed under standard conditions with standard chow diet and 12 h light/dark cycles. 1 × 10¹² particles per mouse of Cre TOP-EVs or NEO-TOP-EVs were administered by tail vein injection (150 μl PBS). After 7 days, mice were sacrificed and analyzed for tdTomato expression.

### 2.11 Immunoffuorescence staining

Tissues were fixed in formalin for 24 h and embedded in paraffin. Paraffin-embedded organs were cut into 5 μm sections, mounted on slides, and dried at 55 °C for 1 h. Sections were deparaffinized and rehydrated (3× Tissue Clear for 5 min, 2× 99% EtOH for 5 min, 2× 96% EtOH for 5 min, 2× 70% EtOH for 5 min, 3× dH₂O for 5 min). Antigen retrieval was performed by submerging slides in sodium citrate buffer (pH 6.0) containing 0.5% Tween-20 and heating in an autoclave. Slides were washed in PBS (3 × 5 min) and blocked with 3% BSA for 1 h at room temperature. tdTomato-positive cells were stained with a primary RFP antibody (1:250, pre-adsorbed, Rockland Immunochemicals) followed by Goat anti-Rabbit IgG Alexa Fluor 555 secondary antibody (1:200, A-21428, Invitrogen). Nuclei were counterstained with Hoechst (1:10,000, H3570, Invitrogen). Slides were mounted with Fluoromount-G (0100-01, SouthernBiotech) and imaged using a BX53 microscope (Olympus).

Quantification of tdTomato-positive area was performed using ImageJ (NIH). For each mouse, 3 random fields of view were captured per liver section and converted to RGB. The tdTomato-positive area was determined by applying a consistent color threshold across all images. Results are expressed as the percentage of tdTomato-positive area relative to the total image area per field of view.

### 2.12 Cell proliferation and viability assay (CCK-8)

Cell viability was assessed using the Cell Counting Kit-8 (CCK-8, 96992-100TESTS-F, Sigma-Aldrich) assay according to the manufacturer’s instructions. Briefly, cells were seeded into 96-well plates at a density of 10,000 cells/well and incubated in a humidified incubator at 37 °C with 5% CO₂. After treatment, 10 μL of CCK-8 reagent was added directly to each well. Plates were then incubated for 1 h at 37 °C until an obvious color change was observed. Absorbance was measured at 450 nm using a microplate reader. Cell viability was calculated based on the calibration curve.

### 2.13 LDH cytotoxicity assay

Cytotoxicity was quantified by measuring lactate dehydrogenase (LDH) release into the culture supernatant using the CyQUANT™ LDH Cytotoxicity Assay – Fluorescence Kit (Cat. No. C20302, Thermo Fisher Scientific) according to the manufacturer’s instructions. Briefly, on the day of measurement, lysis buffer and stop solution were equilibrated to room temperature, and the LDH reagent stock solution was prepared by combining the supplied Reporter Mix with the Reagent Mix and protecting the mixture from light.

To define assay controls, background wells containing medium only were included. Spontaneous LDH release controls (untreated cells) and maximum LDH release controls were prepared by adding 10 µl of 10x Lysis Buffer to designated wells followed by incubation at 37 °C for 45 min. Culture supernatants (50 µl) were transferred to a separate flat-bottom 96-well plate, mixed with 50 µl of LDH reagent stock solution, and incubated for 10 min at room temperature protected from light. The reaction was terminated by adding 50 µl of Stop Solution, and fluorescence was measured within 1–2 h using a microplate reader (excitation 560 nm, emission 590 nm).

Fluorescence values were background-corrected by subtracting the medium-only signal. Percent cytotoxicity was calculated as:

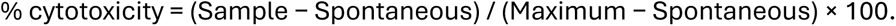

### 2.14 Sanger sequencing

Genomic DNA was extracted from treated Huh7 cells and the PCSK9 target region was amplified by PCR using the following primers: Forward: 5’-acttcagctcctgcacagtcctc-3’, Reverse: 5’-gctccccactacccgtcctc-3’. PCR amplicons were submitted to Macrogen Europe for Sanger sequencing. Base editing efficiency was estimated from Sanger chromatograms using EditR and expressed as the percentage of A•T-to-G•C conversion at the target position.

### 2.15 RT-qPCR

Total RNA was extracted from Huh7 cells using TRIzol reagent (15596026, Invitrogen) according to the manufacturer’s instructions. Complementary DNA was synthesized using the qScript cDNA Synthesis Kit (QuantaBio) according to the manufacturer’s instructions. Quantitative PCR was performed using iQ SYBR Green Supermix (1708885, Bio-Rad) on a CFX Opus 96 Real-Time PCR System (Bio-Rad). PCSK9 mRNA expression was quantified using four primer pairs (Forward-1: 5’-TTGCGTTCCGAGGAGGACGG-3’, Reverse-1: 5’-GCGAGAGGTGGGTCTCCTCC-3’; Forward-2: 5’-GACACCAGCATACAGAGTGACC-3’, Reverse-2: 5’-GTGCCATGACTGTCACACTTGC-3’; Forward-3: 5’-TTGGTGCCTCCAGCGACTGC-3’, Reverse-3: 5’-ACTCGGCCAGGGTGAGCTCC-3’; Forward-4: 5’-TCCACGCTTCCTGCTGCCAT-3’, Reverse-4: 5’-CAGCCAGTCAGGGTCCAGCC-3’) and normalized to human GAPDH (Forward: 5’-GTCTCCTCTGACTTCAACAGCG-3’, Reverse: 5’-ACCACCCTGTTGCTGTAGCCAA-3’). Results are expressed as the mean of four primer pairs relative to the negative control.

### 2.16 PCSKS Elisa

Secreted PCSK9 protein levels were quantified in cell culture supernatants using the Human PCSK9 Quantikine ELISA Kit (DPC900, R&D Systems) according to the manufacturer’s instructions. Cell culture supernatants were collected 3 days after EV treatment, cleared by centrifugation at 500 × g for 5 min, and diluted 20-fold in Calibrator Diluent RD5P prior to analysis. Absorbance was measured at 450 nm with correction at 570 nm using a Multiskan FC microplate reader (Thermo Scientific). PCSK9 concentrations were calculated from a standard curve ranging from 0.625 to 40 ng/mL using a four-parameter logistic curve fit.

### 2.17 LDL uptake assay (pHrodo™ Red LDL)

Low-density lipoprotein (LDL) uptake was measured using the Image-iT™ Low Density Lipoprotein Uptake Kit, pHrodo™ Red (Invitrogen, Cat. No. I34360) according to the manufacturer’s instructions. Cells were seeded one day prior to the assay and, where indicated, serum-starved overnight in assay medium supplemented with BSA to reduce basal lipoprotein uptake.

On the day of the assay, cells were pre-incubated for 30 min at 37 °C with either unlabeled LDL (as a receptor-blocking control) or heparin (to chelate LDL in solution and prevent binding/uptake), followed by incubation with pHrodo™ Red–labeled LDL. pHrodo™ Red LDL was applied at the indicated concentration (typically 10 µg/mL) and incubated at 37 °C for up to 3 h. Because pHrodo™ dyes are minimally fluorescent at neutral pH and fluoresce upon endocytosis into acidic compartments, pHrodo™ Red LDL uptake was quantified in live cells without extensive washing.

Fluorescence was acquired using an RFP setting (approximate excitation/emission maxima 560/585 nm) on EVOS.

### 2.18 Statistical test

Statistical analyses were performed using GraphPad Prism (v.9.3). Data are presented as mean ± s.d. from biological replicates or technical replicates, as indicated in the figure legends. Comparisons between two groups were performed using a two-tailed unpaired t-test with Welch’s correction. Comparisons among multiple groups were analyzed using one-way ANOVA followed by Tukey’s or Dunnett’s multiple comparisons test. P < 0.05 was considered statistically significant.

## 3. Results

### 3.1 Targeting PI(4,5)P₂-rich plasma membrane microdomains enhances cargo loading and delivery efficiency

In the original TOP-EVs platform, protein cargos are recruited to membranes through a chemically induced dimerization system, in which cargo proteins fused to DmrC are conditionally tethered to membrane-anchoring modules fused to DmrA ^34^. In the first-generation design, membrane association was achieved via an N-terminal myristoylation motif on DmrA, enabling EV loading and functional protein delivery ^21^. To improve spatial control over cargo recruitment during EV biogenesis, we next focused on engineering the membrane-targeting step of the TOP-EVs system.

To systematically evaluate whether PI(4,5)P₂ targeting enhances EV cargo loading, we assembled a panel of 14 membrane-interaction modules spanning the major mechanistic modes of PI(4,5)P₂ recognition ^27^(2.1a-2.14a; summarized in Table 1; Fig. 1a,b). These included: (i) structured PI(4,5)P₂-binding domains, such as the Annexin-A2 binding core ^35^, DOC2B-C2 domain ^36^, Epsin1-ENTH domain ^37^, Ezrin-FERM domain ^38^, PI3KC2α-PX domain ^39^, PLCδ1-PH domain ^40^, Tubby protein homolog-Tubby domain ^41^, AP180-ANTH domain ^42^, and the PDZ domains of Syntenin-1 ^43^, which scaffold syndecan-containing membrane microdomains, as well as the adaptor domain Shc1-PID domain, which exhibits membrane association through interactions with acidic phospholipids ^44^; (ii) a PI(4,5)P₂-binding domain that also induces membrane curvature through higher-order self-assembly and multivalent engagement of PI(4,5)P₂, the IRSp53-IBAR domain ^45^; and (iii) polybasic and partially hydrophobic clusters from MARCKS ^46^ and HIV-1 Gag ^47^, which associate with PI(4,5)P₂-rich membranes mainly through electrostatic interactions between positively charged residues and the negatively charged PI(4,5)P₂ headgroups. (iv) a phosphatidylserine-binding module (MFGE8 C2 domain) ^48^, included as a non–PI(4,5)P₂ lipid-binding control.

Each PI(4,5)P₂-associated module was fused to DmrA within the membrane-anchoring construct, replacing the original N-myristoylation motif in the TOP-EVs chemically induced dimerization system. Cre recombinase was expressed as a DmrC fusion, as in the original TOP-EVs design ^21^, and used as the test cargo to quantify loading and EV-mediated functional delivery.

When equal numbers of EVs (normalized by NTA) were analyzed, Ver 2.9a (I-BAR) showed a statistically significant increase in cargo incorporation per particle relative to Ver 1.0, while several other variants showed upward trends that did not reach statistical significance under the current experimental conditions (Fig. 1c,d). In addition, VSV-G and TSG101 levels varied across constructs incorporating different PI(4,5)P₂-binding domains (Fig. 1c), suggesting that PI(4,5)P₂-binding domains influence both cargo incorporation and EV-associated protein composition in a context-dependent manner, rather than functioning as purely passive loading tags.

To assess functional protein delivery, we employed a Cre-loxP reporter system in which successful EV-mediated Cre delivery leads to irreversible recombination and permanent fluorescent reporter activation (Fig. 1e). Functionally, Ver 2.4a (FERM), Ver 2.5a (PH), and Ver 2.9a (I-BAR) produced significantly stronger Cre reporter activation than Ver 1.0 when tested at equivalent particle numbers, as shown by representative fluorescence images (Fig. 1f) and quantified by flow cytometry (Fig. 1g), with Ver 2.9a also exhibiting the most pronounced increase in cargo incorporation per particle. These results are consistent with improved cargo incorporation and more efficient EV-mediated delivery under particle-normalized conditions. Together, these findings indicate that PI(4,5)P₂-guided membrane targeting represents a promising design principle for EV engineering. The observed correlation between PI(4,5)P₂-domain incorporation and enhanced cargo loading is consistent with a model in which lipid-domain targeting improves cargo enrichment, although direct visualization of construct localization to PI(4,5)P₂-enriched membrane domains was not performed in this study.

### 3.2 Recruiting ESCRT-associated modules promotes cargo encapsulation and functional delivery

Although PI(4,5)P₂-guided membrane targeting improved cargo loading (Ver 2.Xa), the original TOP-EVs platform relied solely on N-myristoylation for membrane association and does not explicitly engage late envelopment steps such as membrane neck constriction and scission ^18^. We therefore asked whether reinforcing these later steps through ESCRT-interacting elements could enhance EV budding and cargo encapsulation even in the context of the original N-myristoylated anchor.

To isolate the contribution of ESCRT engagement, we generated a second series of NEO-TOP-EVs (Ver 2.1b-2.7b; summarized in Table 2) by extending the N-myristoylated DmrA anchor with a diverse set of ESCRT-interacting modules derived from viral L-domains ^49^, L-domain–like host factors involved in microvesicle or intraluminal vesicle formation, the canonical ESCRT-recruitment module from cytokinetic abscission machinery, and a Ca²⁺-dependent ESCRT adaptor (Fig. 2a,b). While the cargo module (Cre–DmrC) remained identical to that in TOP-EVs ^21^ (Fig. 2a,b), ensuring that Ver 2.Xb variants specifically report how ESCRT-interacting motifs modify budding and loading on an N-Myr background, without changes in PI(4,5)P₂ targeting.

**Fig. 2.**
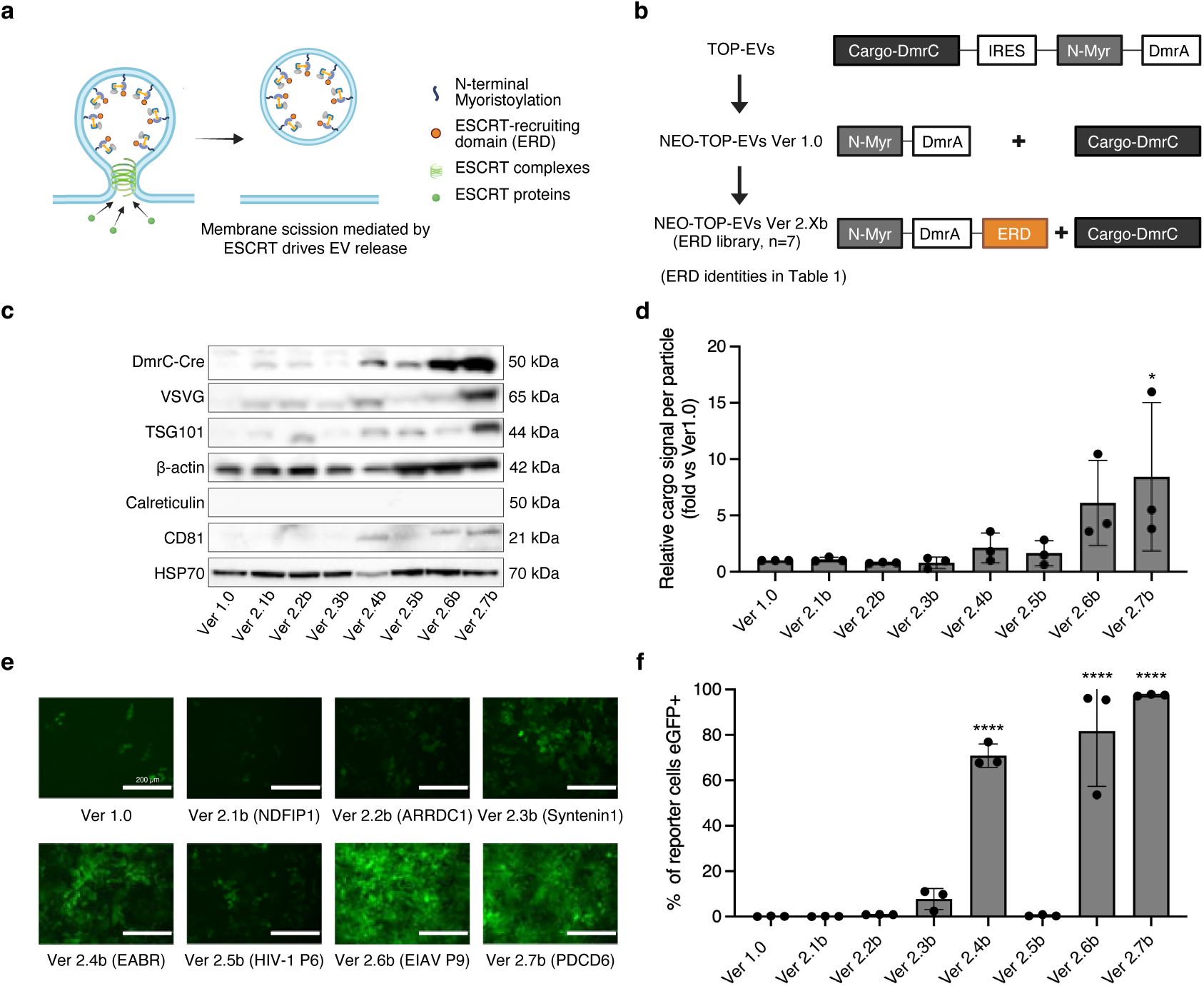
Recruiting ESCRT-associated modules promotes cargo encapsulation and functional delivery. **a**, Conceptual schematic illustrating ESCRT-mediated membrane scission during endogenous EV biogenesis. Recruitment of ESCRT complexes to membrane buds drives membrane constriction and scission, resulting in the release of extracellular vesicles. In the NEO-TOP-EVs platform, this process is engineered through the incorporation of an ESCRT-recruiting domain (ERD) to actively promote vesicle budding and release. **b**, Evolution of the NEO-TOP-EVs architecture incorporating ESCRT-recruiting domains (ERDs). Second-generation constructs (Ver 2.Xb) retain the N-myristoylation anchor while incorporating ERDs to promote membrane scission. **c**, Immunoblot analysis of purified EVs produced by Ver 1.0 and ESCRT-enabled NEO-TOP-EV variants (Ver 2.1b–Ver 2.7b). Blots were probed for DmrC-Cre cargo, vesicular stomatitis virus glycoprotein (VSVG), the EV marker TSG101, β-actin, and EV-associated markers Calreticulin, CD81 and HSP70. **d**, Quantification of cargo incorporation per particle derived from panel c. Densitometry values were normalized to Ver 1.0 control within each blot. Data represent mean ± s.d. from n = 3 independent EV preparations. **e**, Representative fluorescence microscopy images of T47D Cre-reporter cells treated with EVs from Ver 1.0 and Ver 2.Xb constructs. Scale bars, 200 μm. **f**, Flow cytometry quantification of GFP-positive reporter cells corresponding to panel e. Data represent mean ± s.d. from n = 3 independent experiments. For functional delivery assays, 25,000 reporter cells were treated with 1 × 10^11^ EV particles per condition, normalized by NTA. For all bar graphs, statistical comparisons versus Ver 1.0 were performed using one-way ANOVA with Dunnett’s multiple comparisons test. * p < 0.05, **** p < 0.0001.

Although ESCRT-interacting modules have been implicated in promoting vesicle budding in multiple biological systems ^18^, their introduction into NEO-TOP-EVs produced only mild, non-significant upward trends in EV particle counts (Supplementary Fig. S1), suggesting that ESCRT engagement alone is not sufficient to drive consistent increases in vesicle output within this engineered context. Western blot analysis of EVs normalized to equal particle numbers revealed increased cargo incorporation per particle in select Ver 2.Xb variants, with Ver 2.7b (PDCD6) reaching statistical significance (Fig. 2c,d); other variants showed upward trends that did not reach significance, likely reflecting variability across independent preparations. The increased abundance of the ESCRT-associated protein TSG101 in several constructs is consistent with altered engagement of ESCRT-related pathways, although direct recruitment was not assessed. While this observation does not constitute direct evidence of ESCRT recruitment, it is consistent with altered incorporation of ESCRT-associated components during vesicle formation ^3, 18^.

Functionally, ESCRT-enhanced NEO-TOP-EVs (Ver 2.4b-EABR, Ver 2.6b-EIAV P9 and Ver 2.7b-PDCD6) generated significantly stronger Cre reporter activation compared with Ver 1.0 when tested at equivalent particle numbers (Fig. 2e,f), while other Ver 2.Xb variants showed minimal functional improvement, consistent with their modest effects on cargo loading. These observations suggest that select ESCRT-interacting motifs represent an orthogonal handle for improving EV cargo loading and functional delivery, even in the absence of optimized membrane targeting, motivating their combination with PI(4,5)P₂-binding modules in subsequent designs.

### 3.3 Dual PI(4,5)P₂ targeting and ESCRT engagement combinatorially augment EV cargo loading and functional delivery

Building on the observation that PI(4,5)P₂ targeting improves early cargo recruitment, whereas ESCRT-interacting modules enhance late-stage cargo capture, we tested next whether combining both mechanisms would yield cooperative improvements in cargo loading. To this end, we generated a third-generation NEO-TOP-EVs (Ver 3.X) by pairing the three most promising PI(4,5)P₂-binding domains (PH, FERM, and I-BAR) with the three most effective ESCRT-associated regions (EABR, EIAV P9, and PDCD6), producing a 3×3 matrix of dual-envelopment designs (Ver 3.1–3.9, summarized in Table 3). All constructs retained the same DmrA module paired with DmrC-Cre cargo as used in earlier generations, ensuring standardized loading and directly comparable functional assessment.

PI(4,5)P₂–ESCRT combinations produced greater increases in cargo incorporation per particle than either module alone under particle-normalized conditions, as quantified across independent experiments (Fig. 3c,d), with Ver 3.7 and Ver 3.9 showing the most pronounced and statistically significant gains. We note that formal synergy testing (for example, using Bliss independence or highest-single-agent models) was not performed; the term ‘combinatorial’ is used descriptively to indicate that dual-module constructs consistently outperformed either single module alone, without implying a specific mathematical interaction model. These enhancements exceeded the effects observed for the corresponding PI(4,5)P₂-targeting variants alone (Ver 2.Xa), demonstrating that ESCRT engagement provides an additional, independent contribution to cargo encapsulation when layered onto PI(4,5)P₂-guided membrane targeting.

**Fig. 3.**
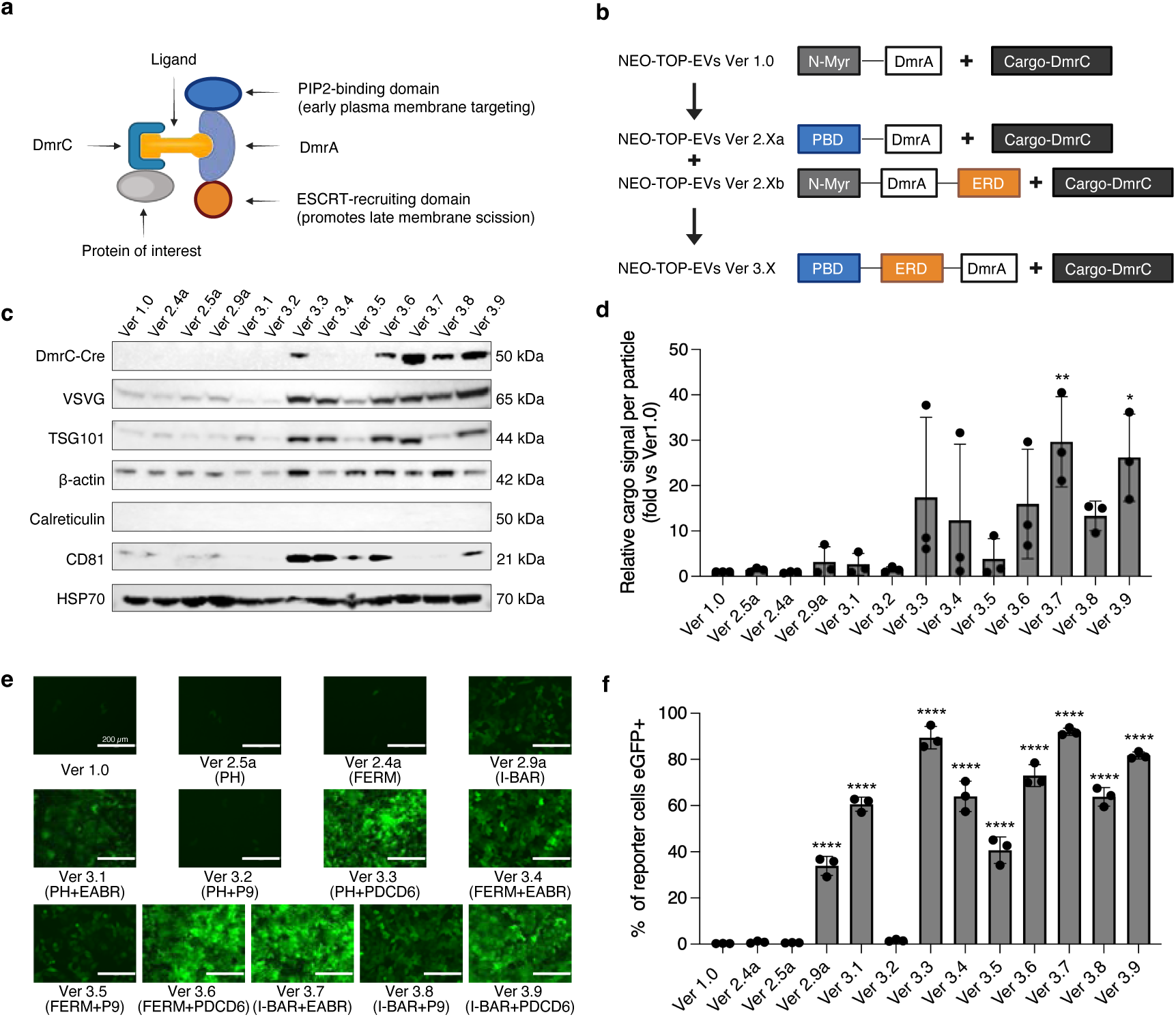
Dual PI(4,5)P₂ targeting and ESCRT engagement combinatorially augment EV cargo loading and functional delivery. **a**, Schematic illustrating the dual-module envelopment design of third-generation NEO-TOP-EVs, in which a PI(4,5)P₂-binding domain (PBD) and an ESCRT-recruiting domain (ERD) are combined within a single PBD– ERD–DmrA fusion protein to coordinate membrane targeting and scission. **b**, Evolution of the NEO-TOP-EVs envelopment architecture from Ver 1.0 through second-generation PI(4,5)P₂-targeting constructs (Ver 2.Xa) and ESCRT-enabled constructs (Ver 2.Xb) to third-generation dual-module constructs (Ver 3.X). **c**, Immunoblot analysis of EVs produced by Ver 1.0, selected Ver 2.Xa constructs, and third-generation NEO-TOP-EVs variants (Ver 3.1–Ver 3.9). Blots were probed for DmrC-Cre cargo, vesicular stomatitis virus glycoprotein (VSVG), the EV marker TSG101, β-actin, and EV-associated markers Calreticulin, CD81 and HSP70. **d**, Quantification of cargo incorporation per particle derived from the immunoblots shown in panel c. Densitometry values were normalized to the Ver 1.0 control within each blot. Data represent mean ± s.d. from n = 3 independent EV preparations. **e**, Representative fluorescence microscopy images of T47D-Cre reporter cells treated with EVs produced by Ver 1.0, second-generation variants (Ver 2.Xa), and third-generation NEO-TOP-EVs constructs (Ver 3.1–Ver 3.9). Scale bars, 200 μm. **f**, Flow cytometry quantification of GFP-positive reporter cells corresponding to panel e. Data represent mean ± s.d. from n = 3 independent experiments. For functional delivery assays, 25,000 reporter cells were treated with 1 × 10^10^ EV particles per condition, normalized by NTA. For all bar graphs, statistical comparisons versus Ver 1.0 were performed using one-way ANOVA with Dunnett’s multiple comparisons test. * p < 0.05, ** p < 0.01, **** p < 0.0001.

These combined effects translated directly into functional delivery. When equal particle numbers were applied to Cre reporter cells, Ver 3.X EVs triggered significantly stronger reporter activation compared with Ver 1.0 and most single-module variants (Fig. 3e,f). Notably, high levels of recombination were achieved at markedly lower particle doses compared to Ver 2.X variants (particle doses detailed in figure legend), reflecting a pronounced increase in per-particle functional potency under particle-normalized conditions. Together, these results demonstrate that coordinated PI(4,5)P₂ targeting and ESCRT engagement generate an envelopment environment that combinatorially enhances both cargo loading and functional delivery, motivating further exploration of additional design features in subsequent generations.

### 3.4 Self-assembly–mediated multivalent clustering amplifies cargo loading and functional delivery in fourth-generation NEO-TOP-EVs

In the third-generation NEO-TOP-EVs library, constructs incorporating an I-BAR domain (Ver 3.7– 3.9) consistently showed the strongest cargo loading and functional delivery. Because the I-BAR domain combines PI(4,5)P₂ binding with intrinsic higher-order assembly, this observation suggested that multivalent clustering of envelopment modules may further improve EV cargo encapsulation.

This observation provided a key conceptual foundation for the fourth-generation NEO-TOP-EVs architecture, in which we sought to generalize and modularize this principle by introducing an engineered self-assembling peptide (SAP) as an independent clustering scaffold. As summarized in Fig. 4b, the NEO-TOP-EVs platform evolved from a minimal membrane-anchoring design (Ver 1.0) to a dual-module PI(4,5)P₂–ESCRT scaffold (Ver 3.X), and finally to a fourth-generation architecture, incorporating an SAP to drive higher-order multivalent clustering.

**Fig. 4.**
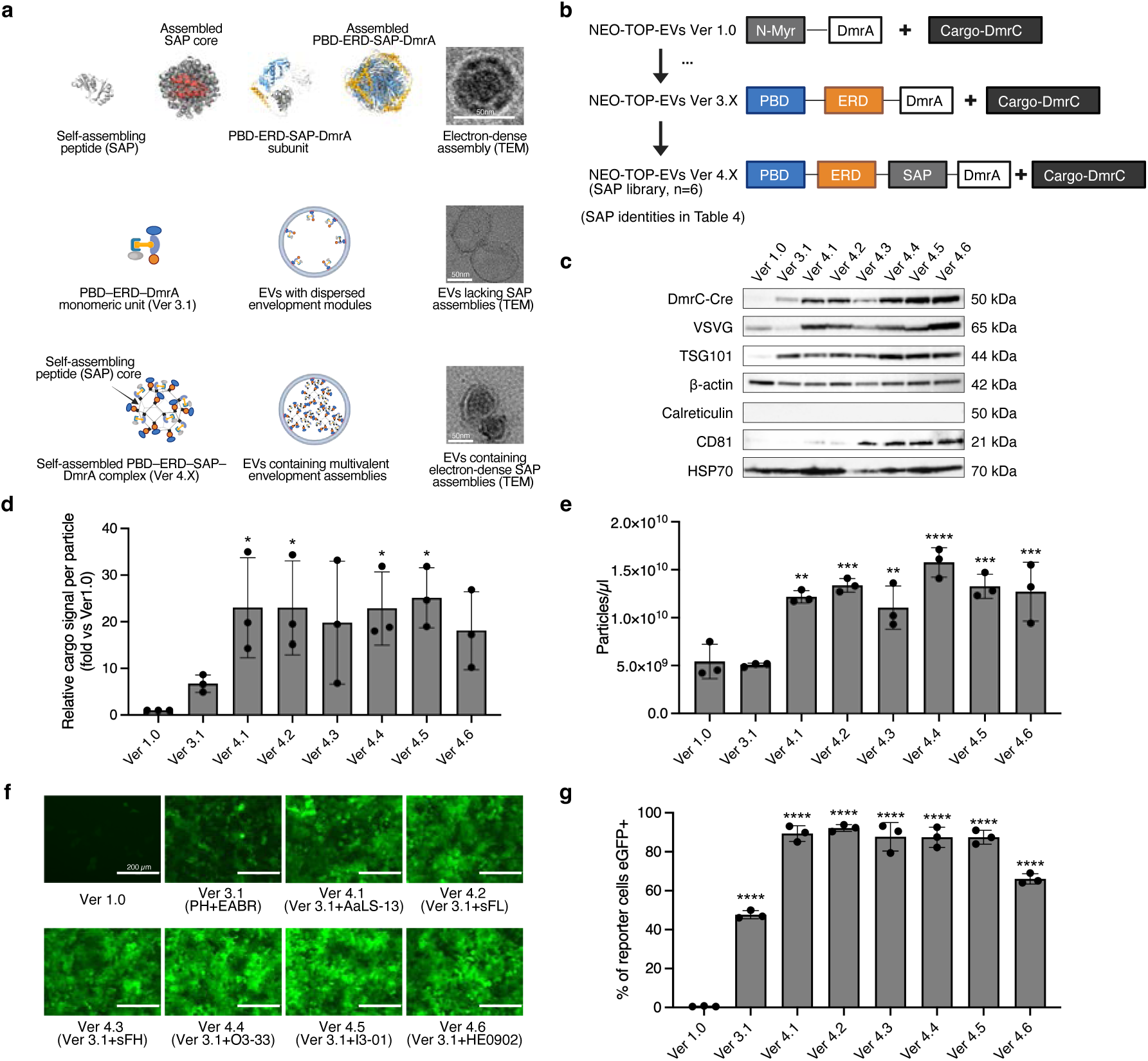
Self-assembly–mediated multivalent clustering amplifies cargo loading and functional delivery in fourth-generation NEO-TOP-EVs. **a**, Structural modeling and transmission electron microscopy illustrating the architecture of self-assembling peptide (SAP) scaffolds used in fourth-generation NEO-TOP-EVs. The SAP forms an oligomeric core that organizes multiple PBD–ERD–SAP–DmrA subunits into higher-order assemblies. Transmission electron microscopy reveals electron-dense structures consistent with SAP-derived assemblies within EVs, whereas EVs lacking SAP modules show dispersed envelopment modules. **b**, Evolution of the NEO-TOP-EVs architecture from the original TOP-EV system to fourth-generation constructs incorporating SAP-mediated multivalent clustering. First-generation constructs (Ver 1.0) employ N-myristoylation–based membrane anchoring. Third-generation constructs (Ver 3.X) combine PI(4,5)P₂-binding domains (PBDs) with ESCRT-recruiting domains (ERDs). Fourth-generation NEO-TOP-EVs (Ver 4.X) introduce SAP modules to promote higher-order assembly of envelopment modules. **c**, Immunoblot analysis of purified EVs produced by Ver 1.0, Ver 3.1 and fourth-generation constructs (Ver 4.1–Ver 4.6). Blots were probed for DmrC-Cre cargo, VSVG, the EV marker TSG101, β-actin, and EV-associated proteins Calreticulin, CD81 and HSP70. **d**, Quantification of cargo incorporation per particle derived from the immunoblots shown in panel c. Densitometry values were normalized to the Ver 1.0 control within each blot. Data represent mean ± s.d. from n = 3 independent EV preparations. **e**, NTA quantification of EV particle production from producer cells expressing Ver 1.0, Ver 3.1 and Ver 4.X constructs. SAP-containing constructs exhibit increased EV particle output relative to earlier generations. Data represent mean ± s.d. from n = 3 independent experiments. **f**, Representative fluorescence microscopy images of T47D Cre-reporter cells treated with EVs produced by Ver 1.0, Ver 3.1 and Ver 4.X constructs, showing enhanced GFP activation following SAP-mediated multivalent clustering. Scale bars, 200 μm. **g**, Flow cytometry quantification of GFP-positive reporter cells corresponding to panel f. Data represent mean ± s.d. from n = 3 independent experiments. For functional delivery assays, 25,000 reporter cells were treated with 5 × 109 EV particles per condition, normalized by NTA. For all bar graphs, statistical comparisons versus Ver 1.0 were performed using one-way ANOVA with Dunnett’s multiple comparisons test. * p < 0.05, ** p < 0.01, *** p < 0.001, **** p < 0.0001.

As an initial strategy, SAP motifs were appended to several top-performing Ver 3.X constructs. Unexpectedly, some of these triple-module designs showed reduced Cre loading compared with their parental Ver 3.X constructs (Supplementary Fig. S2), indicating that enforced clustering is not universally beneficial and may depend on the architecture of the underlying scaffold.

To systematically evaluate SAP integration on a defined background, we selected Ver 3.1 as a well-defined baseline construct for SAP integration. Ver 3.1 is a strong-performing dual-module construct whose PH and EABR components are monomeric in their isolated forms, thereby enabling a controlled evaluation of SAP-mediated clustering effects without confounding intrinsic oligomerization. Six distinct SAP cores, ranging from bacterial-derived, human-derived, and computationally designed scaffolds—with diverse oligomeric architectures and surface-exposed termini were screened (Ver 4.1–4.6, summarized in Table 4). Each SAP was inserted into a fixed N-to-C orientation (PBD–ERD–SAP–DmrA) to preserve consistent module arrangement.

Because SAPs drive higher-order oligomerization, each assembly incorporates multiple copies of the PBD–ERD–DmrA unit, thereby effectively increasing the local density and spatial organization of PI(4,5)P₂-binding domains, ESCRT-recruiting motifs, and cargo-tethering modules (DmrA). This spatial multivalent clustering provides a mechanistic basis for amplified envelopment. Consistent with this design, 3D structural modeling and transmission electron microscopy observations were consistent with the formation of higher-order assemblies of the PBD–ERD– SAP–DmrA fusion proteins (Fig. 4a). In these oligomeric assemblies, the SAP forms a compact central core, while multiple copies of the surface-exposed N-terminal PBD–ERD modules and C-terminal DmrA moieties are arranged peripherally, generating a ring-like outer layer. Representative TEM images further revealed electron-dense substructures in Ver 4.X vesicles that were absent in Ver 1.0. However, systematic quantification of assembly frequency across vesicle populations was not performed.

All Ver 4.X constructs exhibited significantly increased Cre incorporation per particle relative to Ver 3.1 and earlier generations when normalized to equal particle numbers (Fig. 4c,d), demonstrating a pronounced increase in per-particle cargo loading. In parallel, all Ver 4.X constructs exhibited significantly increased EV particle output compared with Ver 1.0 (Fig. 4e), whereas Ver 3.1 did not differ significantly from Ver 1.0 in particle yield. This may suggest that SAP-mediated multivalent clustering of envelopment modules amplifies ESCRT engagement sufficiently to enhance vesicle biogenesis, an effect not achieved by dual-module constructs alone.

Consistent with the increased cargo incorporation per particle, fluorescence imaging revealed markedly enhanced GFP expression in Ver 4.X–treated reporter cells compared with Ver 3.1 and Ver 1.0 (Fig. 4f). Quantitatively, Ver 4.X EVs achieved robust reporter recombination at a particle dose substantially lower than that required for earlier-generation constructs, as detailed in the figure legend (Fig. 4g). Together, these results show that incorporating a modular self-assembly module onto a PI(4,5)P₂–ESCRT scaffold creates a multivalent envelopment architecture that significantly enhances EV biogenesis, cargo loading, and functional delivery efficiency.

### 3.5 The NEO-TOP-EVs envelopment architecture enhances cargo loading independently of VSV-G pseudotyping

A key question for interpreting the NEO-TOP-EVs platform is whether the observed improvements in cargo loading and functional delivery are intrinsic to the engineered envelopment architecture or are contingent upon VSV-G co-expression. To address this, we performed a systematic side-by-side comparison in which selected NEO-TOP-EVs constructs—representing third-generation (Ver 3.7, I-BAR+EABR; Ver 3.9, I-BAR+PDCD6) and fourth-generation (Ver 4.5, I3-01 SAP) designs—were produced with and without VSV-G pseudotyping, alongside first-generation TOP-EVs (Ver 1.0) as a control (Supplementary Fig. S3).

NTA showed that VSV-G co-expression increased overall EV particle yield across all tested constructs (Supplementary Fig. S3a), consistent with the known role of VSV-G in promoting vesicle budding. Immunoblot analysis of particle-normalized EVs revealed that the enhanced cargo loading hierarchy observed across NEO-TOP-EVs generations was preserved in the absence of VSV-G (Supplementary Fig. S3b). Specifically, DmrC–Cre signal was markedly higher in Ver 3.7, Ver 3.9, and Ver 4.5 EVs compared with Ver 1.0, regardless of whether VSV-G was co-transfected. This finding suggests that the biogenesis-inspired envelopment modules—PI(4,5)P₂ targeting, ESCRT engagement, and SAP-mediated clustering—enhance cargo incorporation into EVs through mechanisms that are independent of VSV-G.

In contrast to cargo loading, functional delivery to recipient cells was strictly dependent on VSV-G under the conditions tested. When Cre reporter cells were treated with equal numbers of EVs produced without VSV-G, no detectable reporter activation was observed for any construct, as confirmed by both fluorescence imaging (Supplementary Fig. S3c) and quantitative analysis (Supplementary Fig. S3d). Upon VSV-G pseudotyping, robust reporter activation was restored in a manner that tracked with the envelopment generation: Ver 1.0 produced minimal recombination, whereas Ver 3.7, Ver 3.9, and Ver 4.5 achieved progressively higher levels of reporter activation (Supplementary Fig. S3d).

These results allow a clear functional deconvolution of the NEO-TOP-EVs platform into two separable components: (i) the envelopment architecture, which governs cargo loading efficiency and is independent of VSV-G; and (ii) VSV-G, which is required for cellular uptake and/or endosomal escape and is essential for functional cytosolic delivery under the conditions tested. The observation that cargo loading is enhanced independently of VSV-G provides evidence that the improvements in EV cargo content observed throughout this study are attributable to the biogenesis-inspired envelopment design rather than to indirect effects of VSV-G on vesicle formation. At the same time, the complete absence of functional delivery without VSV-G underscores that the current platform requires a fusogenic element for productive cargo release, and future work will focus on identifying alternative uptake-promoting strategies—such as tissue-targeting nanobodies or fusogenic peptides—to replace VSV-G and enable targeted, non-viral delivery. However, we note that these results are derived from a representative experiment presented in Supplementary Fig. S3; independent biological replication will be required to confirm these findings.

### 3.6 Proof-of-concept in vivo delivery of functional Cre protein to the liver by fourth-generation NEO-TOP-EVs

To subsequently evaluate whether the enhanced cargo loading and delivery potency of fourth-generation NEO-TOP-EVs translate into functional protein delivery in vivo, we used an Ai9 Cre-reporter mouse model that enables sensitive detection of Cre activity through irreversible reporter recombination ^50^ (Fig. 5a). Cre-reporter mice received a single intravenous injection with equal numbers of TOP-EVs or fourth-generation NEO-TOP-EVs (Ver 4.5, 1 × 10¹² particles per mouse) carrying DmrC–Cre cargo.

**Fig. 5.**
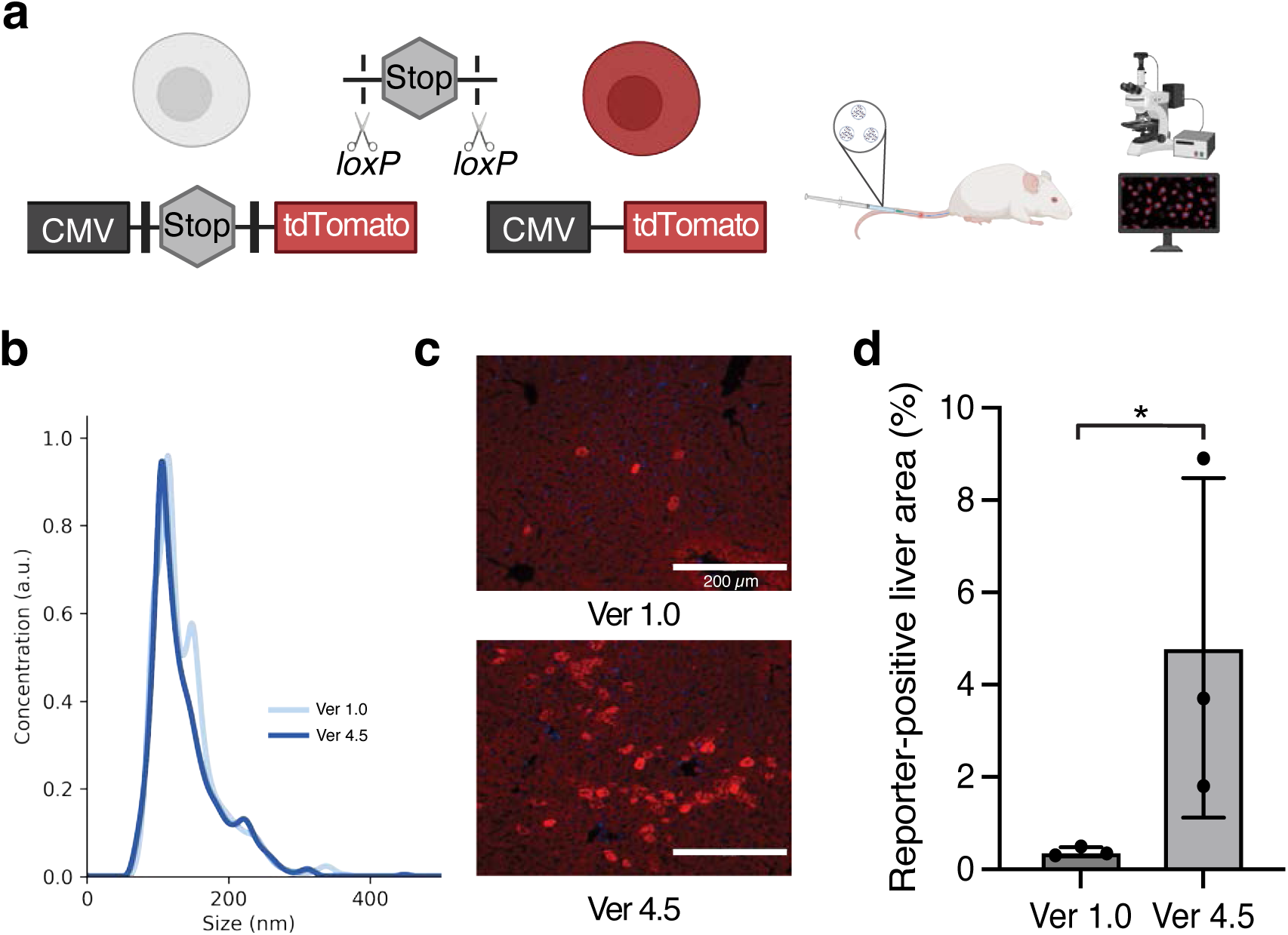
Proof-of-concept in vivo delivery of functional Cre protein to the liver by fourth-generation NEO-TOP-EVs. **a**, Experimental schematic illustrating systemic delivery of Cre-loaded extracellular vesicles in Ai9 Cre-reporter mice. Engineered EVs encapsulating Cre recombinase were administered by tail-vein injection, and Cre-mediated recombination was assessed in target tissues by TdTomato fluorescence activation. **b**, NTA showing size distributions of EVs produced by Ver 1.0 and NEO-TOP-EVs (Ver 4.5) constructs, demonstrating comparable particle size profiles between EV populations. **c**, Representative fluorescence microscopy images of liver sections following systemic administration of Cre-loaded EVs. Robust TdTomato reporter activation is observed in mice treated with Ver 4.5, whereas minimal recombination is detected in Ver 1.0-treated controls. Nuclei were counterstained with Hoechst (blue). Scale bars, 200 μm. **d**, Quantification of TdTomato-positive liver area from 3 random fields of view per liver section following treatment with Ver 1.0 or Ver 4.5, corresponding to panel c. Each dot represents an individual mouse. Bars indicate mean ± s.d. (n = 3 mice per group). Statistical significance was determined by two-tailed unpaired t-test with Welch’s correction, *p < 0.05. For in vivo delivery experiments, mice received 1 × 10¹² EV particles in 150 μL PBS by tail-vein injection. EV doses were normalized to equal particle numbers as determined by NTA.

NTA confirmed comparable size distributions between TOP-EVs and fourth-generation NEO-TOP-EVs (Fig. 5b).

Liver sections harvested after administration of first-generation TOP-EVs exhibited minimal reporter activation, with TdTomato signal largely absent from hepatocytes (Fig. 5c). In contrast, mice receiving fourth-generation NEO-TOP-EVs displayed robust and regionally enriched Cre-mediated reporter recombination within the liver parenchyma, indicating efficient delivery of functional Cre protein in vivo (Fig. 5c). Quantitative analysis of random fields of view from liver sections revealed a significant increase in reporter-positive liver area in mice treated with fourth-generation NEO-TOP-EVs compared with TOP-EVs at matched particle dosing (Fig. 5d), although inter-animal variability was substantial.

We acknowledge several important limitations of this in vivo experiment. The study included only two treatment groups (n = 3 mice per group), a single dose, a single time point, and assessment of a single organ (liver). Furthermore, VSV-G pseudotyping was used, which contributes to liver tropism via LDL receptor-mediated uptake and does not allow the independent contribution of the NEO-TOP-EVs envelopment architecture to tissue targeting to be assessed. Collectively, these results provide preliminary proof-of-concept evidence that the combined engineering advances in fourth-generation NEO-TOP-EVs can support functional protein delivery to the liver in vivo, though the independent contributions of the envelopment architecture and VSV-G-mediated tropism cannot be distinguished under the current experimental design. Building on this in vivo validation, we next evaluated whether NEO-TOP-EVs could support delivery of larger and more complex genome-editing cargoes.

### 3.7 NEO-TOP-EVs enable efficient and low-toxicity delivery of CasG RNPs

To further evaluate the use of the NEO-TOP-EVs platform for delivering large and functional genome-editing cargoes, we replaced the Cre recombinase payload with Cas9 RNP complexes and benchmarked NEO-TOP-EVs against a previously reported virus-like particle (VLP) system ^51^. Cas9 was fused to the DmrC domain to enable envelopment into NEO-TOP-EVs via DmrA–DmrC heterodimerization, while sgRNA was co-expressed from a separate U6-driven plasmid co-transfected into producer cells to assemble active Cas9 RNPs prior to vesicle packaging.

Immunoblot analysis confirmed efficient incorporation of Cas9 into NEO-TOP-EVs, with Cas9 readily detected in purified vesicles (Fig. 6a,b). In contrast, VLP samples displayed both uncleaved Gag–Cas9 fusion and cleaved Cas9 species, consistent with the design of the VLP system. To enable quantitative assessment of genome-editing activity, we employed a stoplight reporter system ^32^ in which Cas9-mediated indel formation induces a frameshift that activates eGFP expression (Fig. 6c).

**Fig. 6.**
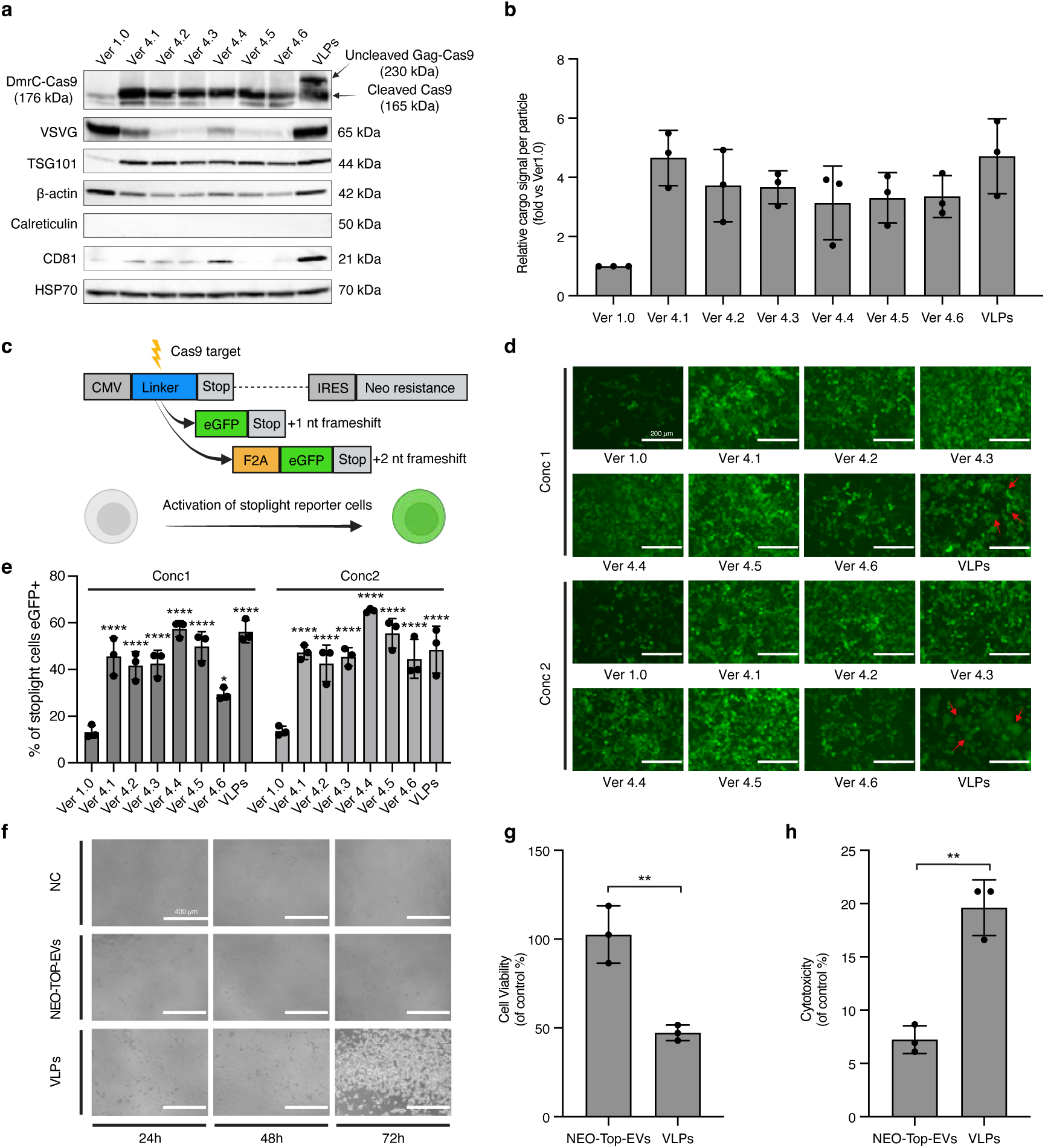
NEO-TOP-EVs enable efficient and reduced-toxicity delivery of Cas9 RNPs. **a**, Immunoblot analysis of purified NEO-TOP-EVs and virus-like particles (VLPs) loaded with Cas9 cargo. Blots were probed for DmrC-Cas9, vesicular stomatitis virus glycoprotein (VSVG), the EV marker TSG101, β-actin, and EV-associated proteins Calreticulin, CD81 and HSP70. In NEO-TOP-EV samples, Cas9 was detected as a single cleaved species. In VLP samples, both uncleaved Gag–Cas9 fusion and cleaved Cas9 were detected, consistent with the design of the VLP system. **b**, Quantification of Cas9 cargo incorporation per particle derived from the immunoblots shown in panel a. Densitometry values were normalized to the Ver 1.0 control within each blot. Data represent mean ± s.d. from n = 3 independent EV preparations. **c**, Schematic of the Cas9 stoplight reporter system used to quantify genome-editing activity following vesicle-mediated Cas9 RNP delivery. Cas9-induced indel formation restores the reading frame and activates eGFP expression in reporter cells. **d**, Representative fluorescence microscopy images of stoplight reporter cells treated with Cas9-loaded NEO-TOP-EVs (Ver 4.1–Ver 4.6) or VLPs at two particle doses. Robust GFP activation indicates efficient genome editing. Scale bars, 200 μm. **e**, Flow cytometry quantification of genome-editing efficiency corresponding to panel d, expressed as the percentage of eGFP-positive reporter cells. No significant difference was detected between NEO-TOP-EVs (Ver 4.1–4.6) and VLPs at either concentration. Data represent mean ± s.d. from n = 3 independent experiments. **f**, Representative images of reporter cells treated with Cas9-loaded NEO-TOP-EVs (Ver 4.5) or VLPs and monitored over 24, 48 and 72 hours. **g**, Quantification of cell viability in Huh7 cells measured using the CCK-8 assay following treatment with NEO-TOP-EVs (Ver 4.5) or VLPs. Data are presented as percentage of control. **h**, Quantification of cytotoxicity in Huh7 cells measured by lactate dehydrogenase (LDH) release following treatment with NEO-TOP-EVs (Ver 4.5) or VLPs. Data are presented as percentage of control. For functional delivery assays (panels d and e), 10,000 Cas9 stoplight reporter cells per well were seeded prior to EV treatment. EVs were applied at two particle doses (Conc1 = 5 × 10¹⁰; Conc2 = 1 × 10¹¹ particles per condition), normalized by nanoparticle tracking analysis (NTA). Data represent mean ± s.d. from n = 3 independent experiments. For cytotoxicity assays (panels f, g, and h), Huh7 cells were seeded at 10,000 cells per well prior to treatment with NEO-TOP-EVs (Ver 4.5) or VLPs at Conc2 (1 × 10¹¹ particles per condition), normalized by NTA. Data represent mean ± s.d. from n = 3 technical replicates of a representative experiment. For e, statistical comparisons versus Ver 1.0 were performed using one-way ANOVA with Dunnett’s multiple comparisons test; NEO-TOP-EVs versus VLPs were compared by unpaired t-test. For g and h, statistical significance was determined by a two-tailed unpaired t-test with Welch’s correction. * p < 0.05, ** p < 0.01, **** p < 0.0001.

NEO-TOP-EVs carrying Cas9 RNPs mediated robust reporter activation across multiple fourth-generation envelopment architectures (Ver 4.1–4.6), reaching editing efficiencies comparable to those achieved by the VLP system at matched particle doses (Fig. 6d,e). At both tested concentrations, NEO-TOP-EVs consistently produced high levels of eGFP-positive reporter cells, demonstrating efficient functional delivery of Cas9 RNPs under particle-normalized conditions.

Notably, while VLPs achieved similarly high editing efficiencies at elevated particle doses, they induced evident cytopathic effects, including extensive cell rounding and detachment within 72 hours of treatment (Fig. 6f). In contrast, NEO-TOP-EVs were well tolerated under the tested condition, even at high particle concentrations. Quantitative viability measurements using CCK-8 assays revealed near-baseline metabolic activity in NEO-TOP-EV–treated cells relative to untreated controls, whereas VLP-treated cells exhibited a marked reduction in viability (Fig. 6g). Consistently, LDH release assays demonstrated significantly higher membrane damage in VLP-treated cells compared with NEO-TOP-EVs (Fig. 6h). We note that these cytotoxicity measurements are derived from technical replicates of a representative experiment; independent biological replication will be required to confirm these findings.

Together, these results demonstrate that NEO-TOP-EVs enable efficient Cas9 RNP delivery with editing performance comparable to VLPs under particle-number–normalized conditions, while exhibiting reduced cytotoxicity in a preliminary assessment; however, cargo-equivalent comparisons will be needed to further characterize the relative safety profiles of these platforms.

### 3.8 NEO-TOP-EVs enable efficient adenine base editor RNP delivery for precise therapeutic genome editing of PCSKG in vitro

To further evaluate the versatility of the NEO-TOP-EVs platform for delivering next-generation genome-editing modalities, we next investigated the delivery of adenine base editors (ABEs), which enable precise A•T-to-G•C conversion without introducing double-strand breaks ^28^. We selected ABE8e, a highly active evolved adenine base editor, as a test case and assessed its delivery and editing activity using both reporter and endogenous disease-relevant targets ^52^.

We first employed a stoplight ABE reporter system in which ABE-mediated base conversion restores the correct reading frame and activates eGFP expression ^33^ (Fig. 7a). NEO-TOP-EVs (Ver 4.5) loaded with ABE8e induced robust activation of reporter cells, with editing efficiencies above 90% across a wide range of particle concentrations (Fig. 7b,c), demonstrating highly efficient functional delivery of ABE8e.

**Fig. 7.**
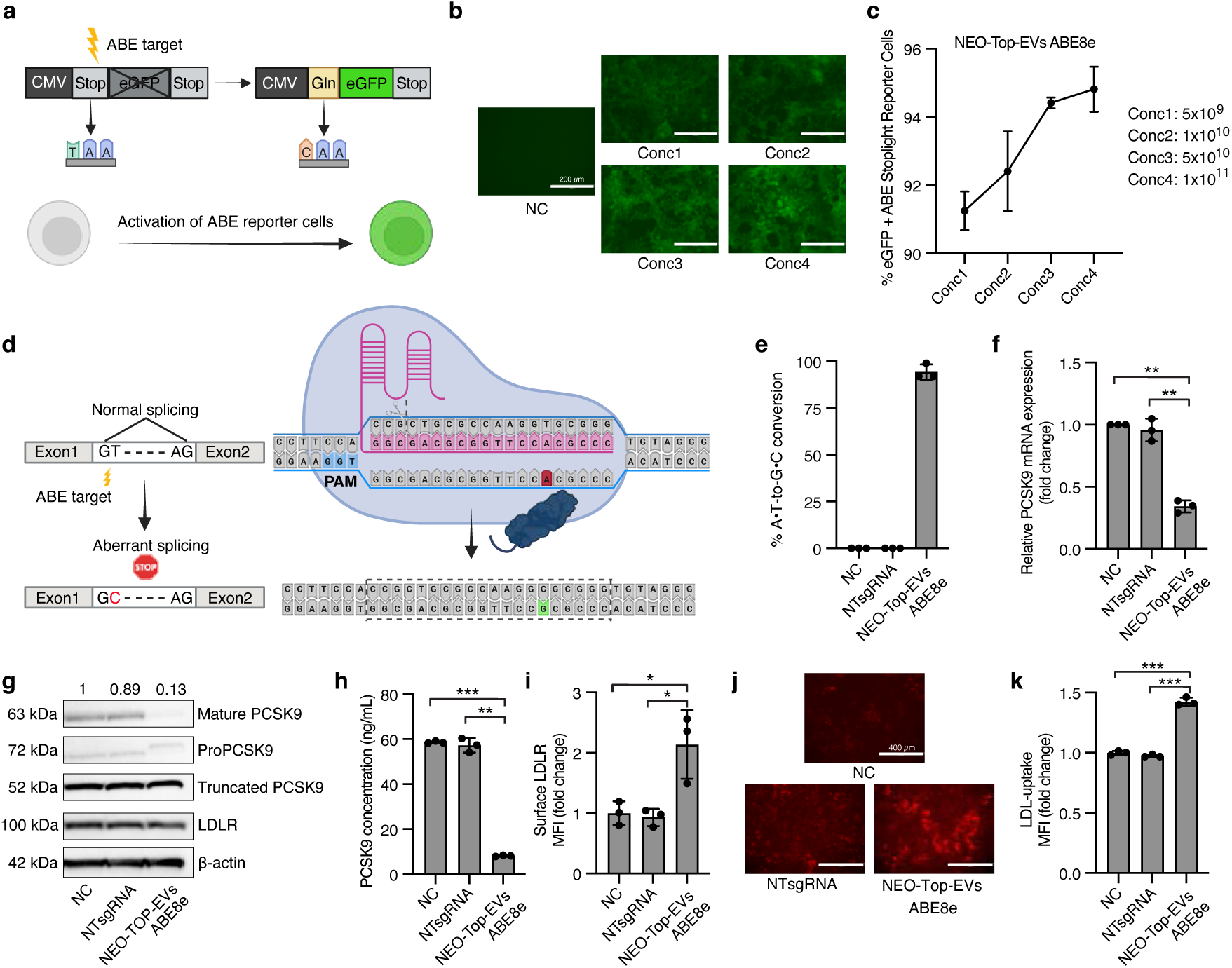
NEO-TOP-EVs enable efficient adenine base editor RNP delivery for precise therapeutic genome editing of PCSK9 in vitro. **a**, Schematic of the adenine base editor (ABE) stoplight reporter system used to quantify base-editing activity following NEO-TOP-EV-mediated delivery of ABE8e. **b**, Representative fluorescence microscopy images of ABE stoplight reporter cells treated with ABE8e-loaded EVs at increasing particle doses (Conc1 = 5 × 10⁹, Conc2 = 1 × 10¹⁰, Conc3 = 5 × 10¹⁰, Conc4 = 1 × 10¹¹ EV particles per condition, normalized by NTA). Scale bars, 200 μm. **c**, Flow cytometry quantification of base-editing efficiency corresponding to panel b, expressed as the percentage of eGFP-positive reporter cells. Note that y-axis is shown from 90% to highlight differences across concentrations; all conditions achieved >90% editing efficiency. **d**, Schematic illustrating ABE-mediated disruption of a canonical splice donor site within PCSK9. **e**, Sanger sequencing analysis of PCSK9 editing at the target splice site, shown as the percentage of A•T-to-G•C conversion. No editing was detected in NC or NTsgRNA controls. **f**, Quantification of PCSK9 mRNA levels by qRT–PCR, normalized to NC (set to 1). **g**, Immunoblot analysis of PCSK9 and LDLR protein levels following ABE8e-mediated splice-site disruption. **h**, Quantification of secreted PCSK9 protein levels in cell supernatants measured by ELISA. **i**, Quantification of cell-surface LDLR expression measured by flow cytometry, shown as mean fluorescence intensity (MFI) fold change relative to NC. **j**, Representative fluorescence microscopy images of DiI-labeled LDL uptake across treatment conditions. Scale bars, 400 μm. **k**, Quantification of LDL uptake measured by flow cytometry, shown as MFI fold change relative to NC. For reporter assays (panels b and c), 10,000 ABE stoplight reporter cells per well were seeded prior to EV treatment. For endogenous PCSK9 editing experiments (panels e–k), 10,000 Huh7 cells per well were seeded prior to treatment under three conditions: negative control (NC), non-targeting sgRNA control (NTsgRNA), and NEO-TOP-EVs loaded with ABE8e at Conc4 (1 × 10¹¹ EV particles), normalized by NTA. All quantitative data for panels e–k are presented as mean ± s.d. from n = 3 technical replicates from a representative experiment. Statistical significance was determined by one-way ANOVA with Tukey’s multiple comparisons test. *p < 0.05, **p < 0.01, ***p < 0.001; ns, not significant.

To validate precise base editing at an endogenous locus, we targeted a canonical splice donor site within the PCSK9 gene. This splice-site targeting strategy for PCSK9 has been previously established as an effective approach for gene disruption using adenine base editors ^30, 51^. ABE-mediated A-to-G conversion at the GT splice donor site disrupts normal splicing, leading to aberrant transcript processing and generation of a truncated PCSK9 protein (Fig. 7d). Sanger sequencing analysis indicated highly efficient base conversion, with an estimated ∼90% A•T-to-G•C conversion at the target site following NEO-TOP-EVs–mediated ABE8e delivery, whereas no detectable editing was observed in control conditions (Fig. 7e).

Consistent with efficient splice-site disruption, NEO-TOP-EV–mediated delivery of ABE8e resulted in a marked reduction in PCSK9 mRNA levels in Huh7 cells (∼65% decrease relative to NC), whereas cells treated with a non-targeting sgRNA showed no significant change (Fig. 7f). This was accompanied by a corresponding reduction in mature PCSK9 protein levels (Fig. 7g). Functionally, disruption of PCSK9 resulted in a pronounced decrease in secreted PCSK9 levels (Fig. 7h), accompanied by increased cell-surface LDL receptor (LDLR) expression (Fig. 7i) and enhanced LDL uptake capacity (Fig. 7j,k).

We note that genome-wide off-target editing analysis was not performed in this study. While PCSK9 splice-site targeting using ABE8e has been previously validated with established guide RNA designs ^30^, comprehensive off-target profiling (for example, GUIDE-seq or CIRCLE-seq) will be essential prior to any translational advancement of this approach.

Together, these results demonstrate that NEO-TOP-EVs enable efficient ABE8e delivery and support precise editing at a therapeutically relevant endogenous locus, providing a foundation for future in vivo evaluation.

## 4. Discussion

In this study, we present NEO-TOP-EVs as a modular extracellular vesicle platform inspired by the endogenous principles of EV biogenesis. By integrating PI(4,5)P₂-mediated membrane targeting, ESCRT machinery-driven budding, and controlled self-assembly into a modular envelopment architecture, NEO-TOP-EVs enable efficient and low-toxicity delivery of diverse macromolecular cargoes, including Cas9 ribonucleoproteins and adenine base editors. Rather than focusing on cargo loading alone, our work establishes a generalizable framework for leveraging EV biogenesis principles in engineering, thereby shifting the design focus from passive cargo recruitment toward active promotion of vesicle formation.

A core design principle of the NEO-TOP-EVs platform is that efficient EV-mediated protein delivery requires coordination of multiple steps in vesicle biogenesis. In natural EV biogenesis, vesicle budding is not driven by membrane association alone but requires the coordinated recruitment of membrane-binding proteins, curvature-generating scaffolds, and ESCRT machinery to defined membrane microdomains. By abstracting this principle, we coupled PI(4,5)P₂-targeting domains with ESCRT-recruiting motifs and cargo-tethering modules in a stepwise manner. Importantly, the progressive performance gains observed from minimal membrane anchoring (Ver 1.0) to PI(4,5)P₂-guided targeting (Ver 2.X), and finally to combined PI(4,5)P₂–ESCRT engagement (Ver 3.X) are consistent with contributions from the coordinated action of multiple envelopment modules.

Our mechanistic interpretation, however, remains correlative. Although increased TSG101 incorporation and improved functional output are consistent with enhanced engagement of ESCRT-related pathways, direct visualization of ESCRT recruitment and construct localization at budding sites was not performed. Furthermore, producer-cell expression levels of the DmrA fusion constructs were not systematically characterized across the library; differences in expression, stability, or folding across architectures may therefore contribute to the observed loading differences independently of enhanced sorting. In addition, immunoblot densitometry provides only a relative, semi-quantitative measure of cargo abundance, and single-vesicle analytical approaches or quantitative mass spectrometry will be needed for absolute per-particle quantification. Future studies combining quantitative imaging, controlled expression analysis, and single-vesicle characterization will be required to validate the proposed mechanism.

Notwithstanding these mechanistic uncertainties, the performance patterns across construct generations offer several design insights. From a biogenesis-inspired perspective, PI(4,5)P₂ may contribute not only to spatial membrane targeting but also to local membrane remodeling through multivalent protein engagement. In the context of NEO-TOP-EVs, this effect is likely reinforced by curvature-generating modules and ESCRT-associated factors, together promoting vesicle envelopment. As a representative example, analysis of the third-generation NEO-TOP-EVs library revealed that constructs incorporating an I-BAR domain (Ver 3.7–3.9) consistently exhibited superior cargo loading and functional delivery. The I-BAR domain is known to combine strong PI(4,5)P₂ binding with intrinsic self-assembly into higher-order lattices, and the ability to induce and stabilize membrane curvature. The robust performance of these variants across both molecular and functional readouts suggests that membrane curvature generation and multivalent lattice formation are not simply supportive elements but instead play an important role in promoting EV budding. By simultaneously clustering envelopment modules and remodeling local membrane topology, I-BAR–containing constructs likely create a highly favorable physical environment for ESCRT recruitment and vesicle scission. This mechanistic insight motivated the design of the fourth-generation NEO-TOP-EVs architecture, in which self-assembly was abstracted and modularized through the introduction of engineered self-assembling peptides as independent clustering scaffolds. By decoupling self-assembly from membrane-binding and ESCRT-recruiting functions, the SAP-based architecture enables modular control over oligomeric geometry, valency, and spatial organization of envelopment modules.

Interestingly, SAP incorporation did not universally enhance all third-generation constructs. Several PI(4,5)P₂-binding domains (such as FERM and I-BAR) and ESCRT-associated regions (such as PDCD6) possess intrinsic dimeric or multimeric tendencies. Enforced clustering through an additional oligomeric SAP may therefore introduce steric or topological incompatibilities that hinder productive membrane engagement rather than enhance it. This observation suggests an important design constraint: multivalency is beneficial only when properly matched to the native oligomeric state and spatial footprint of the envelopment modules. As such, effective EV engineering requires not maximal clustering, but coordinated and structurally compatible assembly of envelopment modules.

Consistent with this interpretation, Ver 3.1—comprising monomeric PH and EABR modules— provided a well-defined and modular baseline for SAP integration. In this context, SAP-driven self-assembly produced a highly ordered multivalent architecture, in which a compact SAP core organized multiple copies of surface-exposed PBD–ERD modules and cargo-tethering domains into a ring-like arrangement. This higher-order organization simultaneously enhanced per-vesicle cargo loading and EV production yield, demonstrating that spatial clustering of envelopment modules is a key determinant of EV biogenesis efficiency when appropriately engineered.

Collectively, these engineering modifications represent a substantial departure from the biophysical and compositional properties of endogenous EVs. While NEO-TOP-EVs are inspired by and exploit natural EV biogenesis pathways, the incorporation of heterologous membrane-targeting domains, ESCRT-recruiting motifs, and self-assembling scaffolds is likely to alter vesicle composition, surface properties, and potentially immunogenicity and in vivo biodistribution in ways that remain to be systematically characterized. NEO-TOP-EVs should therefore be understood as engineered delivery vehicles that leverage EV biogenesis principles, rather than as faithful mimics of endogenous EVs—a distinction with direct implications for their translational development. In addition, all EVs in this study were produced in HEK293FT cells; whether the NEO-TOP-EVs envelopment architecture performs equivalently in therapeutically relevant producer cell types, and how producer cell identity influences vesicle composition and in vivo behavior, remains to be determined.

Despite these current limitations, NEO-TOP-EVs compare favorably with existing delivery platforms in several key respects. Compared with conventional delivery strategies such as AAV vectors, lipid nanoparticles, and virus-like particles, NEO-TOP-EVs offer a distinct set of advantages by engineering the cellular vesicle production machinery rather than constructing an exogenous carrier. AAV rely on capsid-mediated encapsulation and suffer from limited cargo capacity, pre-existing immunity, and challenges in repeated dosing ^53, 54^. Lipid nanoparticles, while highly effective for nucleic acid delivery, are poorly suited for protein or ribonucleoprotein cargos and often exhibit dose-limiting toxicities ^55^. Virus-like particles enable efficient protein delivery but retain viral structural components and may induce cytopathic effects at high particle doses ^56^. In contrast, NEO-TOP-EVs leverage endogenous EV biogenesis pathways to generate EV-based vesicles that are expected to be free of exogenous nucleic acid templates, with high cargo loading capacity, robust functional delivery, and a potentially favorable safety profile. Direct benchmarking against a VLP-based Cas9 delivery system (Banskota et al., Cell 2022) illustrates this safety distinction in particular: NEO-TOP-EVs achieved comparable editing efficiencies while exhibiting reduced cytotoxicity at elevated particle numbers in a preliminary assessment. We note, however, that normalization was performed by particle number rather than cargo content, and the observed cytotoxicity difference may partly reflect higher cargo content per NEO-TOP-EV particle, differences in VSV-G density, or differences in particle composition rather than an intrinsic safety advantage of the EV chassis. Cargo-normalized benchmarking will be important to establish the relative safety profiles of these platforms. Furthermore, functional delivery in the current system is contingent on VSV-G pseudotyping, which also contributes to liver tropism in vivo and prevents independent assessment of the envelopment architecture’s contribution to cellular entry; replacing VSV-G with tissue-specific targeting moieties will be an important next step. Similarly, the platform relies on chemical induction of DmrA–DmrC dimerization using a rapalog compound during EV production; potential residual ligand in purified EV preparations and immunogenicity of the dimerization domains themselves will require systematic evaluation prior to clinical advancement. It should also be noted that random field-of-view–based quantification may underestimate local delivery efficiency, as reporter-positive cells were not homogeneously distributed throughout the parenchyma; region-of-interest–based or single-cell quantification approaches will therefore be needed for more accurate assessment of in vivo delivery efficiency in future studies.

Beyond Cas9 RNP delivery, NEO-TOP-EVs enable efficient transport of next-generation genome-editing modalities, including adenine base editors. Using ABE8e as a stringent test case, NEO-TOP-EVs support highly efficient base editing in a stoplight reporter system, with editing efficiencies exceeding 90% across a wide range of particle doses. Importantly, this performance translates to precise editing at an endogenous disease-relevant locus. The ability to achieve robust splice-site editing and downstream functional modulation of the PCSK9–LDLR axis via a protein-based, EV-mediated delivery system without reliance on nucleic acid templates distinguishes NEO-TOP-EVs from nucleic acid–dependent approaches and supports their potential for durable therapeutic intervention.

PCSK9 represents a clinically validated therapeutic target for cardiovascular disease, with monoclonal antibodies and siRNA-based therapeutics already approved for lowering LDL cholesterol ^57–59^. Base editing of PCSK9 has been proposed as a potential approach for durable suppression of hepatic PCSK9 expression^30^. Our results demonstrate that NEO-TOP-EVs can deliver ABE8e with sufficient efficiency to induce robust splice-site disruption, suppress PCSK9 secretion, increase cell-surface LDL receptor abundance, and enhance LDL uptake. These findings support further investigation of NEO-TOP-EVs as a promising platform for precision genome-editing therapies targeting metabolic and cardiovascular disorders. We note, however, that genome-wide off-target profiling was not performed in this study, and comprehensive off-target analysis will be essential prior to any translational advancement. Additionally, all quantitative data for endogenous PCSK9 editing experiments (Fig. 7e–k) are derived from three technical replicates of a single representative experiment; independent biological replication will be required for definitive validation.

In summary, the modular architecture of NEO-TOP-EVs—in which membrane-targeting, ESCRT-recruiting, and self-assembly modules can be independently substituted and optimized— provides a flexible foundation for systematic platform development. Key priorities for future work include replacing VSV-G with tissue-specific targeting moieties to enable targeted non-viral delivery, extending the platform to additional cargo types and therapeutically relevant cell populations beyond those tested here, and conducting comprehensive preclinical evaluation including biodistribution, pharmacokinetics, immunogenicity, and genome-wide editing specificity. Together, these advances will be needed to translate the proof-of-concept efficiency demonstrated here into a clinically viable protein-based genome-editing therapeutic.

## Acknowledgements

The authors thank Mark Daniels and Daniek Kapteijn for their experimental support. The authors would like to thank Dr. Linglei Jiang for providing the Cre reporter cell line. The authors would like to thank Dr. Olivier G. de Jong for providing the Cas9 stoplight reporter cell and ABE stoplight reporter cell lines. Schematic illustrations have been created with BioRender.com. During the preparation of this manuscript, the authors used Claude to assist with language editing and proofreading; all content was subsequently reviewed and edited by the authors, who take full responsibility for the published work.

## Author Contributions

S.X. and Q.Y. contributed equally to this work. S.X. conceived the NEO-TOP-EVs platform, designed and performed all experiments including plasmid construct generation, EV production and characterization, reporter assays, and functional delivery studies, and wrote the manuscript. Q.Y. contributed to experimental design and discussion, assisted with plasmid construct generation, and supported EV production. K.Q. assisted with manuscript writing and figure preparation. N.I. contributed to experimental design and provided critical input on the TOP-EVs platform. B.Y. provided scientific input. P.V. provided scientific input and critically reviewed the manuscript. M.A.D.B. performed in vivo experiments and immunofluorescence analysis. C.S.B. assisted with EV production. A.G. provided scientific input. P.A.D. provided scientific input. R.S. provided scientific input and critically reviewed the manuscript. J.X. provided scientific input. Z.L. and J.P.G.S. supervised the project, secured funding, and critically reviewed and edited the manuscript. All authors read and approved the final manuscript.

## Data Availability

The datasets used and/or analyzed in the current study are available from the corresponding authors upon reasonable request.

