## Supplementary figures + Table 1-4 for "An extracellular vesicle biogenesis-inspired engineering platform for efficient protein delivery and therapeutic base editing"

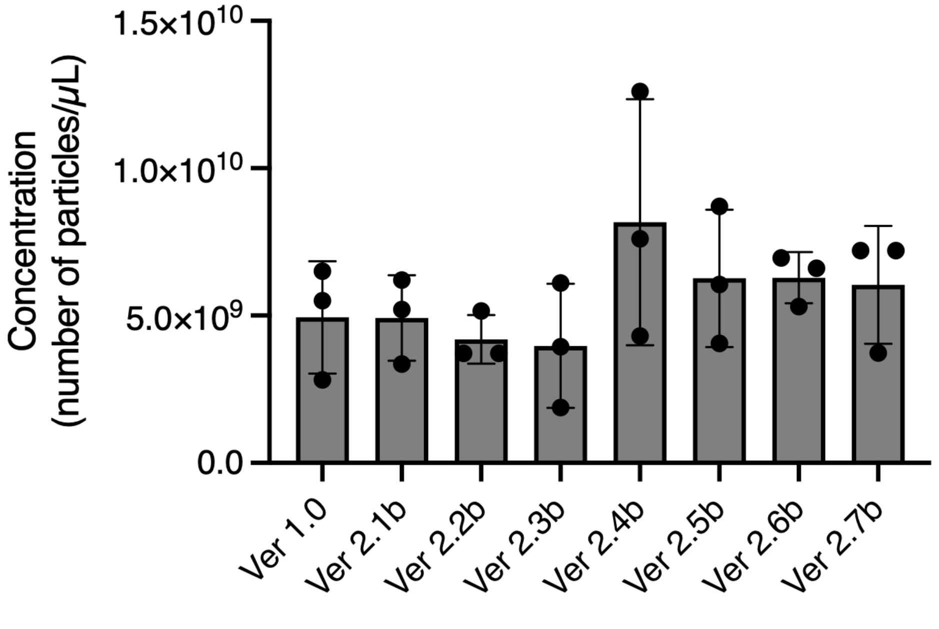


**Supplementary Fig. S1 | Particle production across different Ver 2.Xb constructs.**

Concentration of extracellular vesicle–associated particles produced from different construct versions (Ver 1.0 and Ver 2.1b - 2.7b), as quantified by NTA. Data are presented as particle number per microliter (mean ± s.d., n = 3 independent experiments).

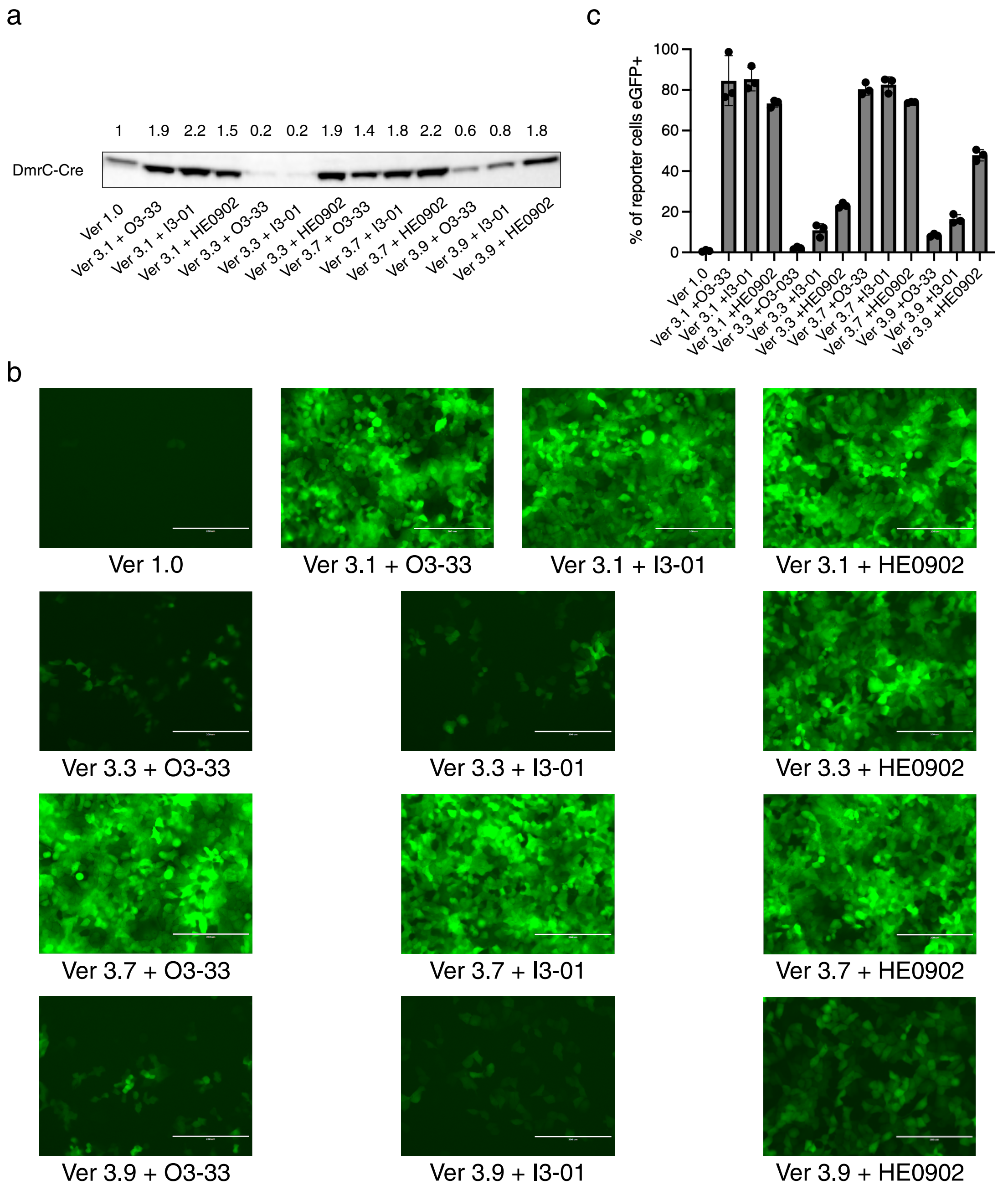


**Supplementary Fig. S2 | Initial integration of self-assembling peptide modules into third-generation constructs does not uniformly enhance cargo loading or functional delivery.**

**a**, Representative Western blot analysis of DmrC–Cre incorporation into extracellular vesicles generated from Ver 3.X constructs with or without appended self-assembling peptide (SAP) modules (O3-33, I3-01, HE0902). Band intensities corresponding to relative Cre incorporation are indicated above the blot. **b**, Representative fluorescence microscopy images of reporter cells following treatment with EVs derived from the indicated constructs, showing GFP activation as a readout of Cre delivery. Scale bars,

200 μm. **c**, Quantification of functional delivery efficiency, expressed as the percentage of eGFP-positive reporter cells following treatment with EVs derived from Ver 3.X constructs with or without SAP modules. Data are presented as mean ± s.d. from a representative experiment. No statistical comparisons were performed. For functional delivery assays, 25,000 reporter cells were seeded prior to EV treatment. EVs were normalized to $5 \times 10^9$ particles per condition as determined by NTA.

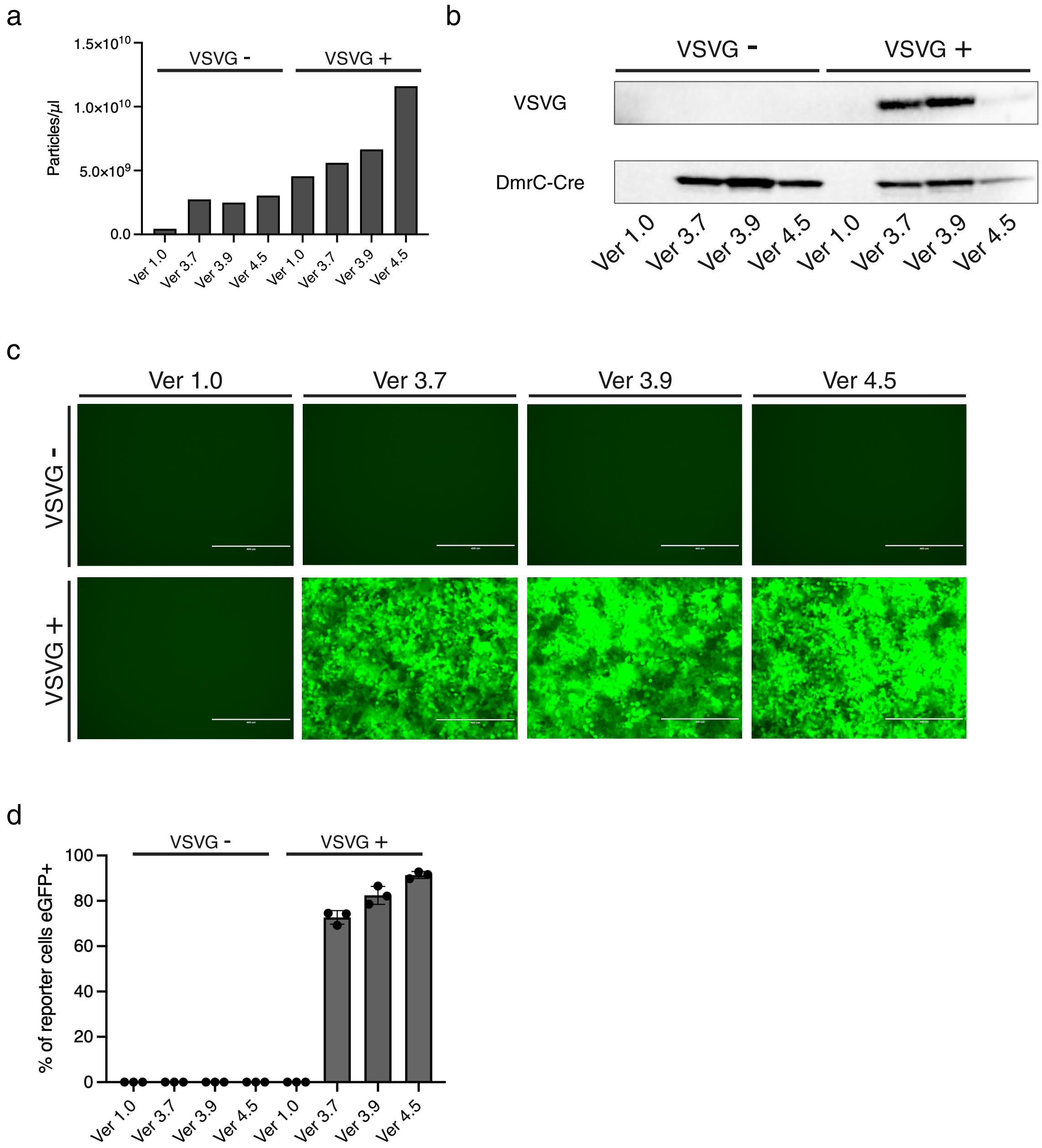


**Supplementary Fig. S3 | The NEO-TOP-EVs envelopment architecture enhances cargo loading independently of VSV-G pseudotyping**

**a**, Quantification of particle production from different construct versions (Ver 1.0, Ver 3.7, Ver 3.9 and Ver 4.5) in the presence or absence of VSVG, measured by NTA. **b**, Western blot analysis of VSVG incorporation and DmrC–Cre expression in vesicle preparations with or without VSVG. **c**, Representative fluorescence microscopy images showing reporter activation (eGFP) in recipient cells following delivery of vesicles with or without VSVG. Scale bars, 200 µm. **d**, Quantification of reporter-positive ($eGFP^+$) cells under the indicated conditions, demonstrating enhanced functional delivery in the presence of VSVG.  Data are presented as mean ± s.d. from a representative experiment. No statistical comparisons were performed. EVs were normalized to $5 \times 10^9$ particles per condition as determined by NTA.

#### Table 1 – PI(4,5)$P_2$ binding domains

| | *Protein* | *AA range* | *Species* | *Effector Domain* | *Version* |
|---|---|---|---|---|---|
| 1. | AP180 | 14-145 | Human | ANTH Domain | Ver 2.1a |
| 2. | Epsin-1 | 12-144 | Human | ENTH Domain | Ver 2.2a |
| 3. | Annexin A2 | 33-336 | Human | Binding Core | Ver 2.3a |
| 4. | Ezrin | 2-296 | Human | FERM Domain | Ver 2.4a |
| 5. | PLCD1 | 21-190 | Human | PH Domain | Ver 2.5a |
| 6. | P3KC2A | 1422-1538 | Human | PX Domain | Ver 2.6a |
| 7. | SHC1 | 156-339 | Human | PID Domain | Ver 2.7a |
| 8. | Syntenin1 | 114-298 | Human | PDZ Domain | Ver 2.8a |
| 9. | IRSp53 | 1-250 | Human | I-BAR Domain | Ver 2.9a |
| 10. | DOC2B | 266-399 | Human | C2 Domain | Ver 2.10a |
| 11. | Tubby protein homolog | 244-506 | Human | Tubby Domain | Ver 2.11a |
| 12. | MFGE8 | 230-387 | Human | C2-Like Domain | Ver 2.12a |
| 13. | MARCKS | 152-176 | Human | N-Myr + a cluster of positively charged residues | Ver 2.13a |
| 14. | Gag | 14-31 | HIV-1 | N-Myr + Highly Basic Region | Ver 2.14a |

### Table 2 – ESCRT-Recruiting domains

| | Protein | AA range | Species | Function | Features | Version |
|---|---|---|---|---|---|---|
| 15. | NDFIP1 | 2-116 | Human | Intraluminal vesicles formation | L-domain-like motifs | Ver 2.1b |
| 16. | ARRDC1 | 285-433 | Human | Microvesicles formation | L-domain-like motifs | Ver 2.2b |
| 17. | Syntenin1 | 2-60 | Human | Intraluminal vesicles formation | L-domain-like motifs | Ver 2.3b |
| 18. | CEP55 | 160-217 | Human | Cytokinesis | EABR region | Ver 2.4b |
| 19. | Gag p6 | 449-497 | HIV-1 | Virus budding | L-domains | Ver 2.5b |
| 20. | Gag p9 | 436-486 | EIAV | Virus budding | L-domains | Ver 2.6b |
| 21. | PDCD6 | 2-191 | Human | Plasma membrane repair | EF-hands | Ver 2.7b |

Table 3 – Combinatorial architecture of NEO-TOP-EVs Ver 3.X

| *PI(4,5)$P_2$-binding domains* | *ESCRT-recruiting domains* | *Version* |
|---|---|---|
| PH (monomer) | EABR (monomer) | Ver 3.1 |
| PH (monomer) | P9 (monomer) | Ver 3.2 |
| PH (monomer) | PDCD6 (dimer) | Ver 3.3 |
| FERM (dimer) | EABR (monomer) | Ver 3.4 |
| FERM (dimer) | P9 (monomer) | Ver 3.5 |
| FERM (dimer) | PDCD6 (dimer) | Ver 3.6 |
| I-BAR (multimer) | EABR (monomer) | Ver 3.7 |
| I-BAR (multimer) | P9 (monomer) | Ver 3.8 |
| I-BAR (multimer) | PDCD6 (dimer) | Ver 3.9 |

#### Table 4 – Combinatorial architecture of NEO-TOP-EVs Ver 4.X

| ***Self-assembling peptide*** | ***Oligomeric size*** | ***Species*** | ***Version*** |
|---|---|---|---|
| AaLS-13 | 360 | Bacterial | Ver 4.1 (based on Ver 3.1) |
| short Ferritin light chain (sFL) | 24 | Human | Ver 4.2 (based on Ver 3.1) |
| short Ferritin heavy chain (sFH) | 24 | Human | Ver 4.3 (based on Ver 3.1) |
| O3-33 | 24 | Computationally designed | Ver 4.4 (based on Ver 3.1) |
| I3-01 | 60 | Computationally designed | Ver 4.5 (based on Ver 3.1) |
| HE0902 | 240 | Computationally designed | Ver 4.6 (based on Ver 3.1) |
